# Determinants of de novo mutations in extended pedigrees of 43 dog breeds

**DOI:** 10.1101/2024.06.04.596747

**Authors:** Shao-Jie Zhang, Jilong Ma, Meritxell Riera, Søren Besenbacher, Julia Niskanen, Noora Salokorpi, Sruthi Hundi, Marjo K Hytönen, Tong Zhou, Gui-Mei Li, Elaine A. Ostrander, Mikkel Heide Schierup, Hannes Lohi, Guo-Dong Wang

## Abstract

Intensive breeding of dogs has had dramatic effects on genetic variants underlying phenotypes. To investigate whether this also affected mutation rates, we deep-sequenced pedigrees from 43 different dog breeds representing 404 trios. We find that the mutation rate is remarkably stable across breeds and is predominantly influenced by variation in parental ages. The effect of paternal age per year on mutation rates is approximately 1.5 times greater in dogs than humans, suggesting that the elevated yearly mutation rate in dogs is only partially attributed to earlier reproduction. While there is no significant effect of breeds on the overall mutation rate, larger breeds accumulate proportionally more mutations earlier in development than small breeds. Interestingly, we find a 2.6 times greater mutation rate in CG Islands (CGIs) compared to the remaining genome in dogs, unlike humans, where there is no difference. Our estimated rate of mutation by recombination in dogs is more than 10 times larger than estimates in humans. We ascribe these to the fact that canids have lost PRDM9-directed recombination and draw away recombination from CGIs. In conclusion, our study sheds light on stability of mutation processes and disparities in mutation accumulation rates reflecting the influence of differences in growth patterns among breeds, and the impact of PRDM9 gene loss on the de novo mutations of canids.

## Main Text

Decreased costs of genome sequencing and improved bioinformatics pipelines^1^ have made it possible, at scale, to identify the set of new mutations that an individual is born with, by sequencing the trio of father, mother and offspring. This has led to mutation rate estimates from many vertebrates^2,3,4,5^, improving phylogenetic dating and providing insights into evolutionary changes to the mutational process across vertebrates. However, only humans^6,7,8,9,10^ and mice^11^ have had a sufficient number of trios sequenced and analysed for intraspecific analysis determinants of mutation rate. Little is known, therefore, about how mutations accumulate over time in the germline of other mammalian species, including dogs.

Dogs constitute a particularly interesting model for studying mutational processes. First, canids are unique among mammals in lacking a functional *PRDM9* ortholog^6^. In other mammals, *PRDM9* recognizes specific sequence motifs and directs the recombination machinery toward these positions, though in some species only weakly^7^. However, without *PRDM9*, canids position recombination events in open chromatin regions, most notably in CG Islands (CGIs)^6, 12^. Given that recombination is mutagenic^8^, the lack of *PRDM9*-directed recombination in dogs should translate into differences in the genomic distribution of germline de novo mutations (DNMs) compared to other mammalian species with a functional *PRDM9* gene, such as humans.

Furthermore, intensive breeding of dogs over the past two hundred years has fostered an impressive array of phenotypic diversity in, for example, body size^9, 10^ and shape^11^, fur type^13, 14^, coat colour^15, 16^ and breed-specific behavioural and disease enrichment^17,18,19,20^. Association studies have identified allelic variants of large phenotypic impact across breeds. Very strong artificial selection by humans may also have affected DNA repair processes and, thus, mutational processes may differ between breeds.

Here we identify de novo mutations in 390 trios from 43 breeds of dogs raised in similar environments in Finland. Our dataset includes unusually large pedigrees (on average 7.48 trios), allowing us to study potential differences in mutational processes among breeds with different phenotypical makeups and life histories. Moreover, by comparing the accumulation of germline DNMs in CGIs with the rest of the genome, we estimate the mutagenic effects of recombination in dogs, and provide an estimate of the time when *PRDM9*-directed recombination was lost in the canid lineage.

## Dog mutation rates shaped by parental ages

We sequenced dog families collected at the dog biobank at the University of Helsinki, Finland, over the past ∼15 years. Genomes were sequenced to an average coverage of 43.3X from 643 dogs (341 females, 302 males) representing 54 multigenerational families and 404 trios, from a total of 43 distinct dog breeds (Supplementary information 1, Supplementary Data 1). We excluded 14 trios with at least one individual displaying an average coverage lower than 24X, thus retaining 390 trios. The pedigrees vary in relationship structure and size, including 37 extended pedigrees average litter size = 2.4), and 81 trios with multiple siblings (mean = 3.6) (Fig. 1, Supplementary Fig. 1). We applied a stringent pipeline to call DNMs for all 404 trios (Methods and Extended Data Fig. 1). We identified 8,312 high-quality autosome DNMs from these 390 trios, with an average of 21.31 DNMs per trio (95% c.i.: 20.14-22.49, Binomial) and a mean callable genome fraction of 96.52% (95% c.i. 96.44-96.56, Bootstrap). Our dataset includes a hypermutated individual with 230 DNMs (ID: FAM007647), which we excluded from all subsequent analyses. We found that 1.51% of DNMs are in coding regions, similar to what has been previously reported in humans (Extended Data Fig. 2). Searching for genes with several mutations, we found enrichment for neurodevelopmental genes in both dogs and humans, but these were generally in non-coding regions, and the biological significance of this observation is unclear (Supplementary information 2).

**Fig. 1.**
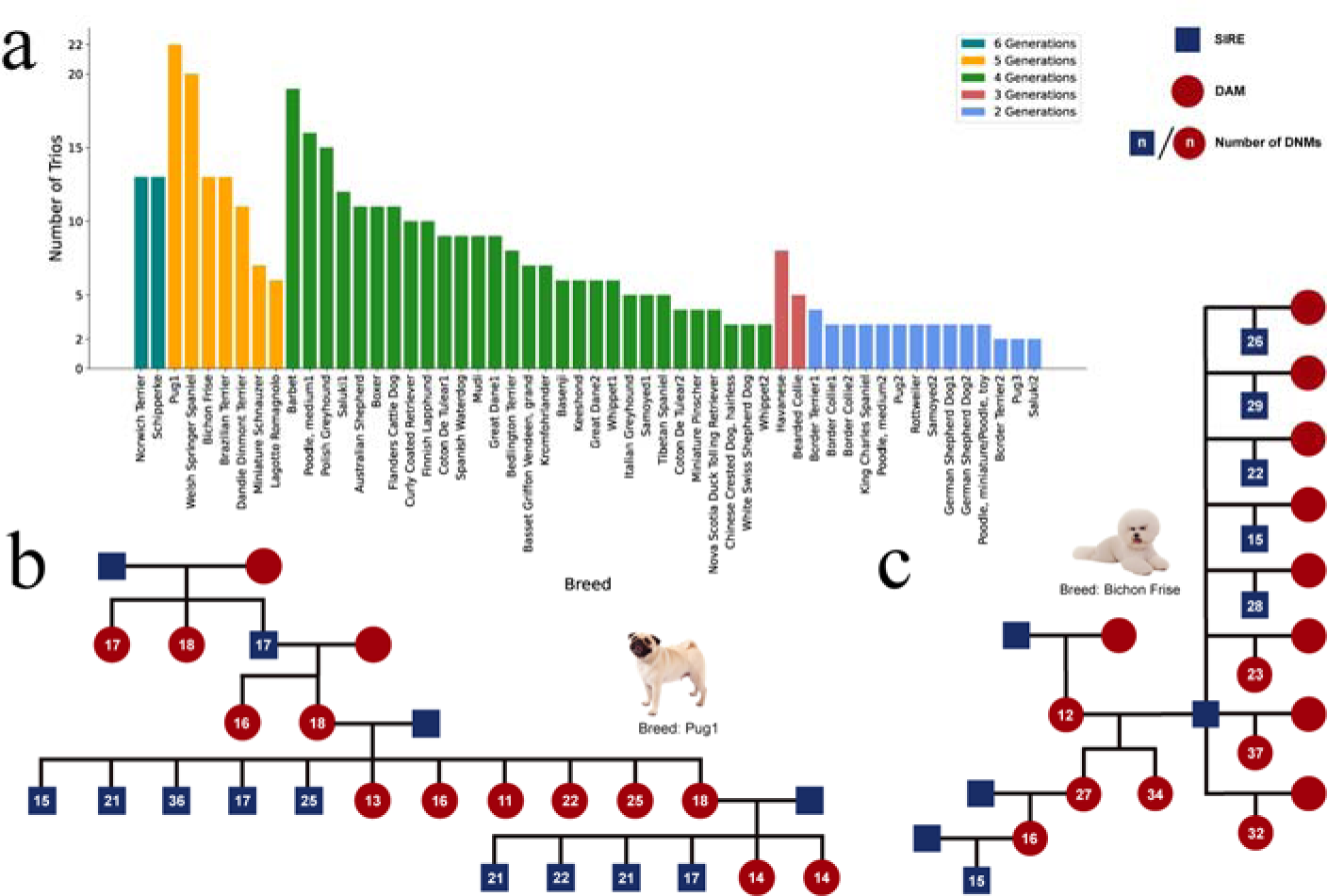
Sample information and statistics of de novo mutations. (a) Number of trios per pedigree. Bar graphs show the corresponding breeds of all 54 pedigrees (horizontal axis), the number of trios in each pedigree (vertical axis), and the number of generations in each pedigree (color of bars). (b) and (c) The demonstration of two representative pedigrees of pug1 and bichon frise, respectively. Blue boxes represent male and red circles represent female, the numbers in them represent the number of DNMs for the individual.

Taking the individual callable genome into account (Methods), we observe an average germline DNM rate of 4.89 × 10^09^ (95% c.i. 4.77 × 10^09^-5.02 × 10^09^, Bootstrap) per base pair, per generation across the pedigrees in autosomes. This estimate is slightly higher than previous pedigree-based estimates in wolves^21^(4.5 × 10^−9^), suggesting a divergence time between dogs and wolves of ∼23,000-30,000 years (Extended Data Table 1). Figure 2a shows per-generation mutation rate estimates from the individual breeds, together with their phylogenetic relationships. The estimated mutation rate per trio differs significantly among breeds (P = 5.4 × 10^−12^, ANOVA). The breed’s effect on the rate per trio is less after accounting for differences in paternal age at conception, but remains statistically significant (P = 0.00014, ANOVA). However, the differences in germline mutation rate across breeds are no longer significant after accounting for parental age differences when we consider rates per litter, instead of treating littermates as independent trios (P = 0.602, ANOVA) (Supplementary information 3).

**Fig. 2.**
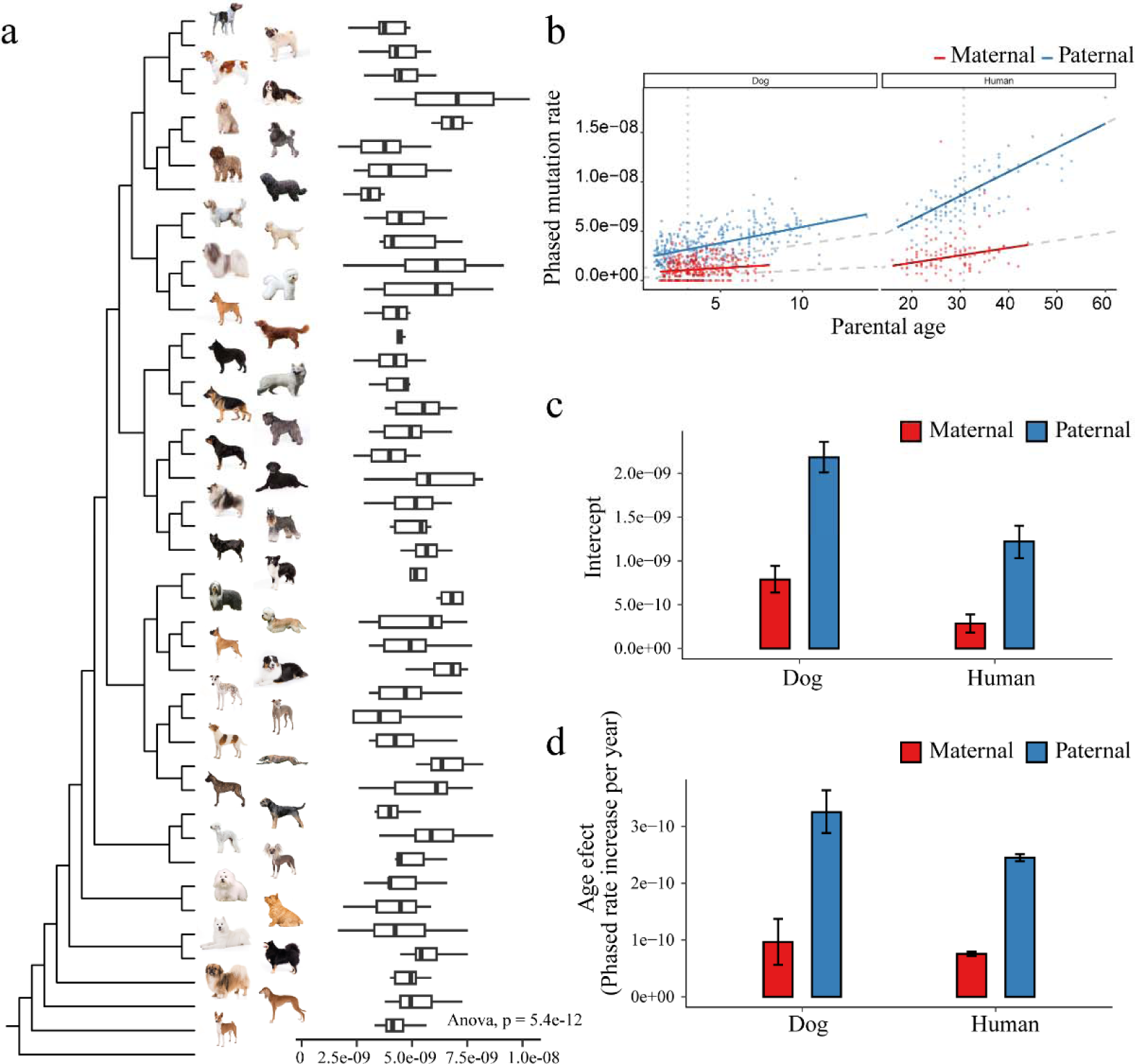
Demonstrations of mutation rates. (a) Mutation rates from the dog breeds together with their phylogenetic relationships. (b) Paternal age and maternal mutation rates in humans and dogs, respectively. (c) Intercept of the mutation rate for male and female in dogs and humans, respectively. (d) Age effect of the mutation rate for male and female in dogs and humans, respectively.

Using read-backed phasing, we determined that the parental origins of 2,586 out of 8,312 DNMs (31.11%). Of the phased DNMs, 75.05% (95% c.i. 73.59 – 76.48, Bootstrap) were of paternal origin, corresponding to a male-to-female mutation ratio of 3.01 (95% c.i. 2.79 – 3.25, Bootstrap). This is less biased than in humans^22^ (0.79% of paternal origin, 3.70), but as for humans, we find a significant association between paternal age and the number of paternal DNMs (Extended Data Fig. 3). We investigated male and female mutation rates as a function of parental ages and compared the dog results to human data^8^. We find that paternal age and maternal age are both significant predictors of mutation rate in dogs (Extended Data Fig. 4) (adjusted R² = 0.3425, P<2 × 10^−16^ and P=0.000118, respectively), as in humans. We also modelled phased mutation rates as a function of parental ages using Bayesian Poisson regression (Figure 2b). We observe significant posterior estimates for paternal age effects on paternal mutation rates (3.25 × 10^−10^, 95% HDI 2.88 × 10^−10^ – 3.63 × 10^−10^) and maternal age effects on maternal mutation rates (9.64 × 10^−11^, 95% HDI 5.61 × 10^−11^ – 1.37 × 10^−10^)(Supplementary information 4). The posterior estimates for paternal intercepts are higher in dogs than humans, suggesting a bigger contribution of mutations accumulated early in development (Figure 2c). Additionally, dogs show a steeper accumulation of paternal mutations per year, with paternal age effect estimates 1.5 times greater than in humans (Figure 2c). This higher yearly accumulation in the male germline, and their shorter generation time. translates into a higher yearly rate of de novo mutation in dogs compared to humans (1.41 × 10^−9^ HDI: 1.37 × 10^−9^ – 1.45 × 10^−9^ vs 3.8 × 10^−10^ HDI: 3.78 × 10^−10^ – 3.82 × 10^−10^).

Paternal age at conception explains less of the variance in paternal germline mutation rates in dogs (McFadden’s R^2^ of 30.47%) than humans (McFadden’s R^2^ of 56.18%), using Poisson regression. McFadden’s R^2^ value is still higher in humans after downsampling the number of DNMs to match that found in dogs (56.06%, Supplementary information 4), suggesting that additional factors may contribute to the variance accumulation of paternal DNMs in dogs. We tested for differences in 21 quantitative phenotypes among breeds including size and lifespan but found that none have a significant effect on the overall mutation rate per litter, based on an ANOVA analysis using parental ages as covariates. (Supplementary information 3).

We next compared the accumulation of germline DNMs with parental age among breeds of dogs of different sizes, assigning each to a category of small, intermediate or large size, based on weight (Figure 3a). We found that large breeds accumulate more paternal DNMs early in development, yielding higher estimates for the intercept (Figure 3b) (3.14 × 10^−9^, HDI: 2.58 × 10^-9^ – 3.7 × 10^−9^) than small breeds (2.03 × 10^−9^, HDI: 1.78 × 10^−9^ – 2.29 × 10^−9^). Conversely, we found that smaller breeds accumulate more paternal mutations per year, as evidenced by a higher paternal age effect on paternal mutation rates in small dogs (3.93 × 10^−10^, HDI: 3.30 × 10^−10^ – 4.55 × 10^−10^) compared to that observed in large dogs (1.66 × 10^−10^, HDI: 7.81 × 10^−11^ – 2.63 × 10^-10^) (Figure 3c). Thus, even though the overall per-generation mutation rate is similar for small and large dogs, we find that the accumulation of germline DNMs through life varies among breeds of differing body sizes. This variation may reflect differences in growth patterns among breeds, with larger breeds having more early cell divisions (faster initial growth) and later puberty^23, 24^, and shorter lifespan^25^.

**Fig. 3.**
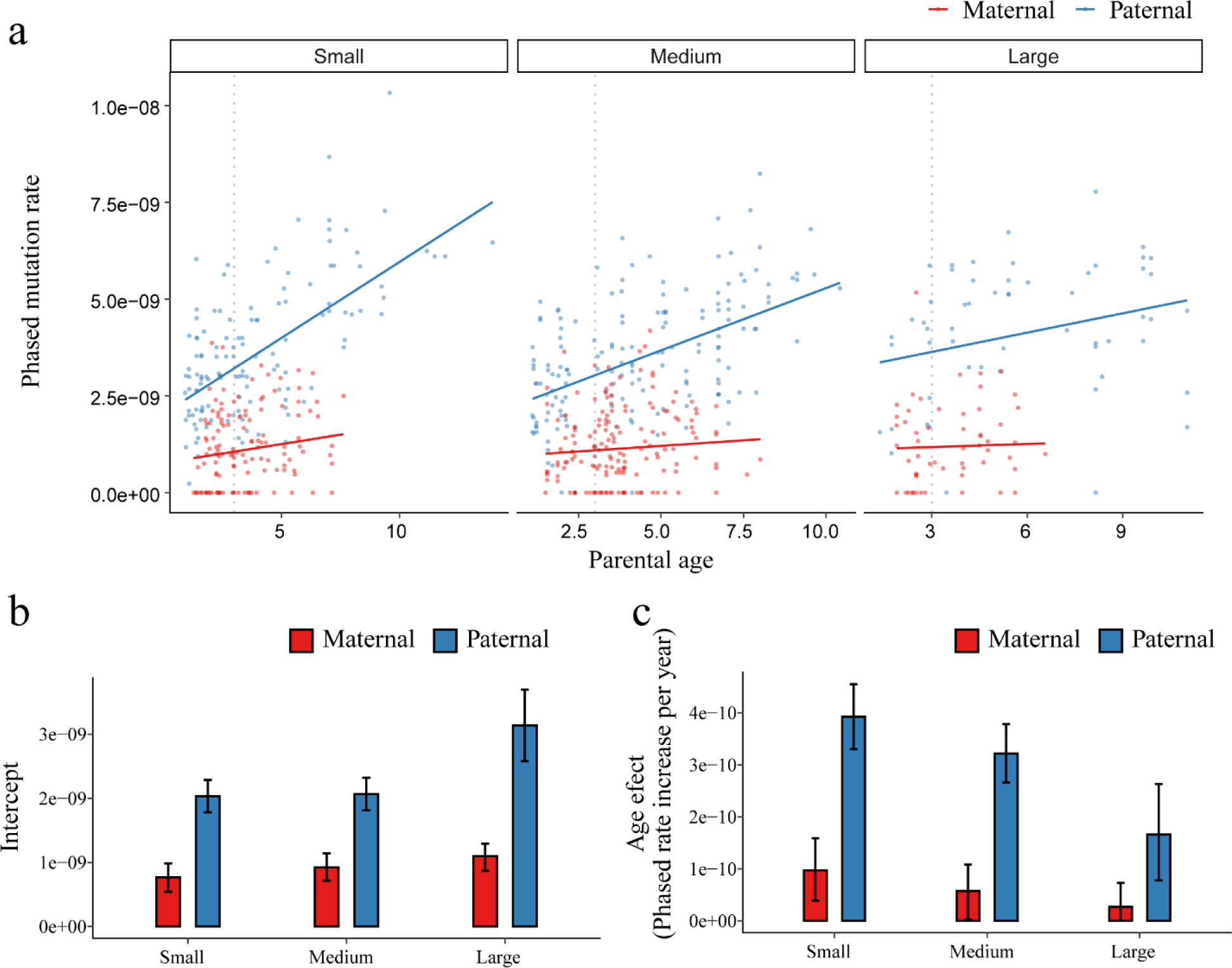
Body sizes contributes to the variance in mutation rates. (a) Phased mutation rates with parental age among small, intermediate, large dog breeds, and humans. (b) The intercept and slope of the mutation rate for paternal and maternal among small, intermediate, large dog breeds. (c) Age effect of the mutation rate for paternal and maternal among small, intermediate, large dog breeds.

## Mutational spectrum in dogs compared to humans

Next, we compared the mutational spectrum of germline DNMs between dogs, mice and humans by stratifying DNMs into eight classes representing the six possible single base pair changes, plus separate categories for C>T mutations in a CpG context, and mutations occurring in CGIs (Methods) (Figure 4a). We find that the spectrum of mutations in dogs is more similar to mice than humans. Notably, dogs show a greater rate of C>T mutations and a smaller rate of T>C mutations than do humans. Intriguingly, this is also the pattern observed when comparing young to old fathers among human pedigrees^7^, suggesting that the mutation spectrum in dogs is more similar to that transmitted by very young parents in humans. In addition, the rate of mutations occurring in CGIs is significantly higher in dogs than in mice and humans.

**Fig. 4.**
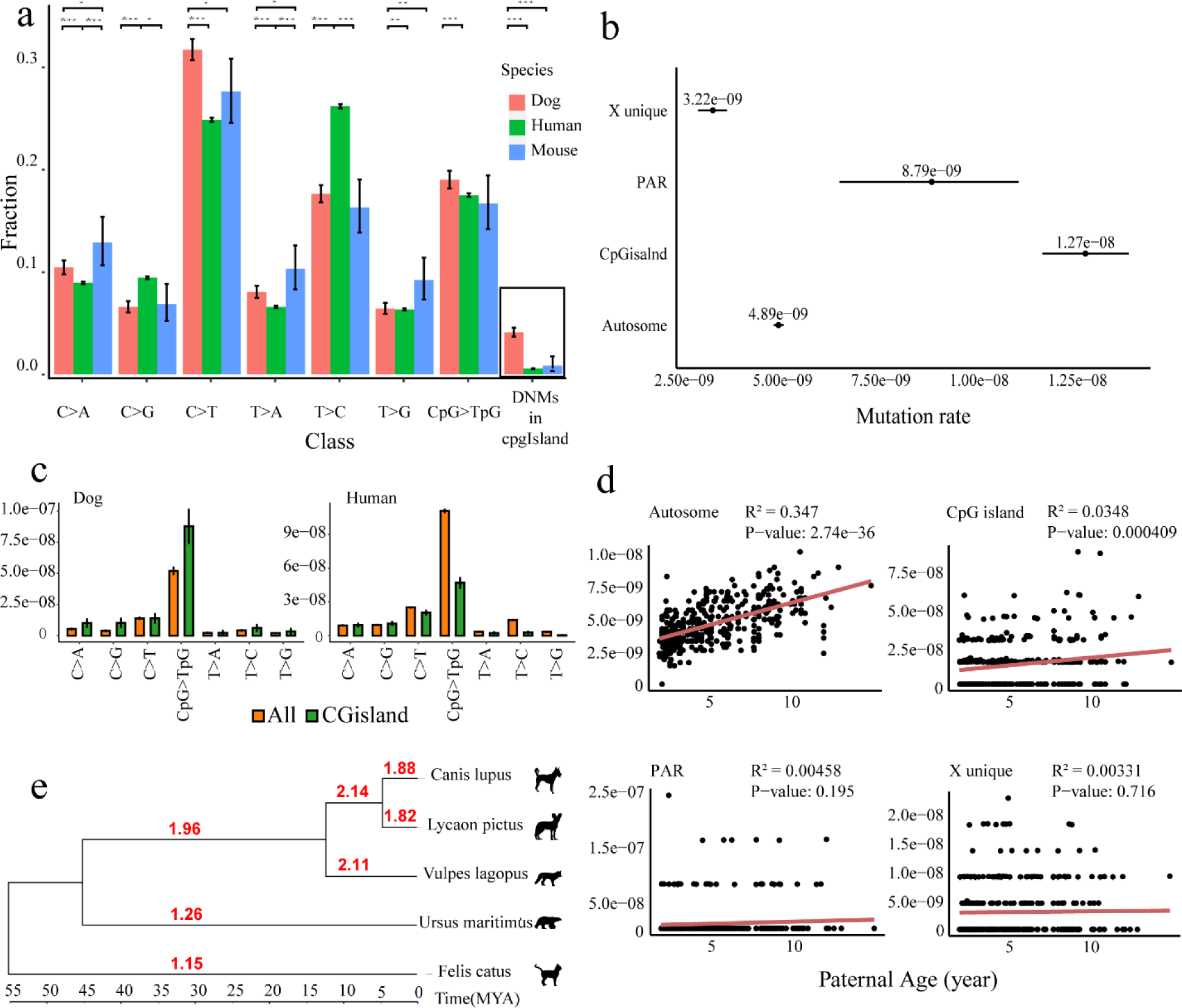
Dog mutational spectrum among species and genomic component. (a) Relative frequency of mutational classes by mouse, dog and human. The black box is a separate class, the number of mutations in CpG islands, and the other seven classes are combined as 1. (b) The comparison of mutation rates in different genomic regions (X chromosomes, PAR regions, CpG islands and Autosomes) of dogs. (c) Comparison of mutation rates of different mutation classes in CpG islands and the whole genome. (d) The mutation rates per individual with parental age of dogs in different genomic regions (X chromosomes, PAR regions, CpG islands and Autosomes). The red lines represent fitted linear regressions only using the paternal age. The black points denote the observed rate. (e) The comparison of evolutionary rate between CpG islands and whole genomes in five species (dogs, African wild Dog, arctic fox, polar bear and domestic cat). The branch length represents the divergent time. The value on each branch is the ratio of evolutionary rate cpg and whole genome evolutionary rate.

These differences in mutational spectrum might be explained by a higher proportion of mutations in dogs occurring earlier in development, i.e., before puberty, as suggested by a significantly larger intercept in the accumulation of DNMs with parental age. To investigate this, we compared the mutational composition of DNMs shared between siblings or half-siblings but found no significant differences between the mutational spectrum in shared mutations and non-shared mutations. We note that this analysis is based on only 79 unique mutations shared by 34 sibling groups, similar to what is observed in a comparable human trio data set^26^ (Extended Data Fig. 5).

## Mutation rate on the X-chromosome

Since the X chromosome spends ⅔ of the time in females but only ⅓ of the time in males, and given that 71.11% of the DNMs are paternal in origin, the mutation rate on the X chromosome is expected to be lower than that of the autosomes. Following our estimated male-to-female mutation rate ratio of 3 (α), we would expect an X-to-autosome mutation rate ratio of 0.83 (95% c.i. 0.82 – 0.84, Bootstrap) ([2(2+α)]/[ 3(1+α)]). We found a lower rate ratio than expecte, however, d (0.66), with an estimated mutation rate on the X of 3.22 × 10^09^ (95% c.i. 2.84 × 10^09^ – 3.58 × 10^09^, Bootstrap) and a mutation rate on the autosomes of 4.89 × 10^09^ (95% c.i. 4.77 × 10^09^-5.02 × 10^09^, Bootstrap) (Figure 4b). Thus, the mutation process on the non-pseudoautosomal regions (PAR) of the X chromosome is slower than on the autosomes. A deviation from this is the PAR of 6.8 Mb^27, 28^ on the rest of the X which is 8.79 × 10^09^ (95% c.i. 6.44 × 10^09^ – 1.10 × 10^08^, Bootstrap), which is 1.8 times higher than the autosomal rate. Since there has to be one recombination event in the PAR in each male meiosis, the PAR should have a recombination rate of approximately 100/6.8=14.7 cM/Mb in males and 1 cM/Mb in females, yielding a sex-averaged recombination rate of 7.85 cM/Mb, which is around eight times higher than the genome average. Given that recombination is mutagenic^8, 29^, this higher rate in the PAR could explain the higher mutation rate observed in this region.

## Recombination shapes the genomic distribution of de novo mutations in dogs

Dogs lack *PRDM9*-directed recombination and are therefore expected to have more recombination events in open chromatin, such as CGIs proximal to genes^7, 29^. Strikingly, dogs display a much higher mutation rate in CGIs than both humans and mice (Figure 4a). This corresponds to a mutation rate of 1.27 × 10^08^ (95% c.i. 1.16 × 10^08^-1.38 × 10^08^, Bootstrap), which corresponds to a 2.6-fold increase in CGIs compared to the rest of the autosomes (95% c.i. 2.37 – 2.82, Bootstrap, Figure 4b). We find that the mutational spectrum of DNMs in CGIs is shifted towards a significantly higher rate after correction for multiple-testing (Bonferroni), of C>A (P = 1.46 × 10^−3^; Binomial test), C>G (P = 9.23 × 10^−6^; Binomial test) and CpG>TpG (3.11 × 10^−9^; Binomial test) in dogs, but not in humans (Figure 4c). As in the case of the PAR data presented above, assuming that recombination in humans is mutagenic^8, 29^, a higher rate of recombination in CGIs could potentially explain the larger number of mutations observed within these regions in dogs. If this is the case, recombination causes 2.00 times more mutations in CGIs compared to the PAR, and the recombination rate should then be expected to be twice as high in the CGIs, i.e., 15.70 cM/Mb. Given that the dog CGIs cover a total of 32 Mb, this would correspond to a recombination length of CGIs of roughly 32 × 15.7=502 cM, implying that about 25.4% of the recombination events occur in CGIs, assuming a total dog map length of 1978 cM^30^. These observations allow a very rough estimation of the mutagenic effect of recombination. In 389 trios we would expect ∼1953 recombination events in CGIs (389 × 5.02). We found a total of 334 mutations in the CGIs, where we expect 334/2.6=128.46 mutations from the autosomes. If this excess of mutations of 334-128.46=205.53 mutations were all due to recombination, this yields an estimate of 0.092 (205.53/2240.64, 95% c.i. 0.08 – 0.10, Bootstrap) mutations per recombination event in CGIs. Our estimated rate of mutation by recombination in dogs is more than 10 times larger than estimates from humans^8, 31^. We speculate that this could be part of the reason for using PRDM9 recombination to draw away recombination from CG Island.

Unlike most mutations accumulated in the germline of males and females from conception to reproduction, the mutational effect of recombination is expected to be independent of parental age (since there is always one round of meiosis). Therefore, we expect mutations in the PAR and in CGIs to be less dependent on parental age than for the rest of the autosomes. Moreover, we also expect a lower paternal age effect on mutations in the X chromosome, given that this chromosome is more influenced by the maternal mutation rate. The correlations shown in Figure 4d are consistent with these assumptions.

## Estimated age of loss of *PRDM9*

Over evolutionary time, following the mutagenic effect of recombination, we would expect a faster rate of evolution in the CGIs of species that lack a functional *PRDM9* gene. We sought to investigate this effect by estimating the ratio of CGIs to autosomal substitution rate along the branches of the phylogeny close to dogs. As expected, this rate is around 2 for canid species that lack a functional *PRDM9* gene, while it is 1.15 and 1.26 in the branches leading to the outgroups Ursus and Felix, which have *PRDM9*-mediated recombination (Figure 4e). Interestingly, we also estimated a rate of around two in the ancestral branch of canids back to the split with Ursus, suggesting that the loss of *PRDM9* must have occurred soon after this split. These estimates would situate the loss of *PRDM9* in canids around 45 million years ago, making it an old evolutionary loss.

## Conclusions

Studying mutations rates in 43 distinct dog breeds demonstrates that the mutation rate is very stable, despite the very strong artificial selection associated with dog breeding. The only life-history trait we found associated with mutation accumulation is breed size, where larger breeds accumulate more mutations early in life, implicating to the negative relationship between weight and lifespan of dog breeds^25^. Whether this is a cause of the shorter life expectancy observed in large breeds is an interesting question for future research.

Male dogs accumulate about 1.5 times more mutations in sperm in their testis per year after puberty compared to humans. The higher yearly mutation rate in dogs compared to humans is, therefore, not only an effect of much shorter generation intervals, but also of a higher intrinsic mutation rate in dog spermatogenesis which is conserved across many different breeds. Interestingly, dogs have also been reported to have a higher somatic mutation rate, which could partly explain the 5-7 times shorter in lifespan observed dogs compared to humans, despite having similar rates of cell divisions per year^32^.

The most conspicuous difference between dogs and humans in the distribution of mutations is with regards to the dogs’ much higher rate in CGIs. We ascribe this to the loss of PRDM9-directed recombination in canids. which then place a large fraction (we very crudely estimate 29%) of the recombinations in the open chromatin associated with CGIs. This implies that these evolve faster in canids, and we could use their rate of evolution together with phylogenetic data to estimate that the loss of PRDM9 occurred prior to canid diversification more than 45 million years ago.

## Method

### Sample collection and information

We collected samples from 643 dogs from 43 breeds, including 54 families, and 404 trios (Supplementary information 1). The dogs were selected from the Finnish dog biobank, and All dogs in the study originate from Finland. In subsequent analyses, we found that when the sequencing depth of parents in a trio is low, it affects the results of DNMs calling. Therefore, we removed 14 trios that included low-depth parental samples (average coverage lower than 24X). As a result, the final number of trios included in our DNMs analysis is 390 (Supplementary information 1).

### Whole Genome Sequencing and Variant Calling of a Large Cohort of Dogs

We used the Covaris system to shear 1-3 μg of DNA into fragments ranging from 200-800 bp. The fragments were then sequenced using the Illumina HiSeq 2000 platform with an average depth of 43.3X. We subsequently used the bwa mem –M algorithm^32^ to map the raw sequence reads to the dog reference genome (Canfam3.1)^34^. We employed PICARD (version 1.96) (https://broadinstitute.github.io/picard/) to remove duplicated reads, and merged BAM files for multiple lanes. The sequences were locally realigned and base-recalibrated using the Genome Analysis Tool Kit (GATK, version 3.7-0-gcfedb67)^35^. To produce the final BAM files, we recalibrated base quality using GATK BQSR. We then used the HaplotypeCaller algorithm in GATK to perform variant calling and generated a gVCF file for each sample. We joint genotyped the gVCF files for each trio to generate a raw VCF file. During the base and variant recalibration, we used a list of known SNPs downloaded from the Ensembl database (ftp://ftp.ensembl.org/pub/release-73/variation/vcf/canis_familiaris/) as the training set. Finally, we filtered the raw VCF files based on the following parameters: “QD < 2.0 || FS > 60.0 || MQ < 40.0 || QUAL < 50.0 || SOR > 3.0 || MQRankSum < –12.5” for further analysis.

### De novo mutations calling

We identified de novo mutations (DNMs) in 404 trios from 54 families using the approach outlined in Supplementary Fig 1 adhered to the guidelines on practices from Bergeron et al^1^. The criteria for DNMs calling using the variant call format (VCF) file of each trio are as follows:

a. The offspring genotype is heterozygous (0/1) and the genotype from the same position from both parents is homozygous (0/0).
b. The mutation must be supported by a maximum of one read in the parents.
c. The genotype quality (GQ) of the DNM is no less than 40 (GQ >= 40).
d. The read depth of any individual in the trio is no less than 12 (min-meanDP = 12), more than half of the average depth of the individual, and not more than twice the average depth (0.5*indDP < DP < 2*indDP)). These depth thresholds are halved for X variants in the chromosomes of male offspring, except for variants in the PAR region.
e. The allelic balance, the fraction of reads supporting the alternative allele), in the child must be greater than 0.25 and less than 0.75. The allelic balance of variants in the X chromosome of male offspring must be greater than 0.75, except for variants in the PAR region.
f. Only single nucleotide mutations are retained.

### De novo mutations filtering

To further remove false positive sites from candidate DNMs, we conducted a filter similar to a manual check with IGV^36^. We used the samtools tool^37^ (samtools tview) to check the reads of all DNMs. This check is based on the bam file which is without realignment. We allowed at most one incorrect read. Incorrect reads refer to reads that differ from the reference genome in the parents and reads that differ from both the reference genome and the DNM in the offspring. Additionally, the offspring’s DNM reads were required to meet the filtering criteria for allele balance, with reads supporting the mutation accounting for 0.25-0.75 of the total number of reads. After excluding unqualified sites, we obtained a final set of 8,565 high-quality DNMs.

### Germline generationally mutation rates

The mutation rate per base pair per generation was estimated as the number of DNMs divided by twice the number of callable sites. The number of callable sites is the number of sites where we would be able to call a de novo mutation in the whole genome. We calculated the number of callable sites for each trio as positions in the genome where parents are homozygous for the reference allele that passed the depth filter applied to DNM calling, i.e no less than 12 and not more than twice the average depth. As in the case of DNM calling, these depth thresholds are halved for X variants in the chromosomes in male offspring, except for variants in the PAR region. Here we need to clarify the terms we used for genome callable site and callable size. We use the “callable site” to refer to the genome position of a haploid genome that passes our quality filters for de novo mutation calling, and we use the “callable size” to refer to the number used as the denominator in mutation rate estimation after considering chromosomal region and individual sex difference. For the X chromosome unique region, male individuals possess only one copy and female individuals will carry two. Thus, the factor for scaling a callable site into callable size is 1 and 2 for male and female respectively. For all other regions, the callable size for each individual will be 2 times of the number of callable sites extracted.

### Mutational classes analysis

We discretize the 12 different single nucleotide mutations into 6 mutational classes (C>A, C>G, C>T, T>A, T>C, and T>G, respectively), differentiating C>T mutations in a CpG context (CpG>TpG) from the rest (C>T). We consider DNMs in CpG islands(CGIs) as a separate class. We obtained the annotation of CGIs from UCSC (https://genome.ucsc.edu/), using the genome assemblies of canFam3 and hg38 for dogs and humans, respectively. For comparison, we also used previously published DNMs from mice (760 DNMs from 40 trios)^9^ and humans (181,258 DNMs from 2,976 trios)^8^.

We assessed the difference in the fraction of DNMs for each mutational class between species using Fisher’s exact test(R Package stats version 4.1.1). For each mutational class, we constructed a 2×2 contingency table by dividing the the DNMs for each species into two categories: belonging to the given mutational class (foreground) and not belonging to the mutational class (background) The resulting P-values were adjusted for multiple testing using Bonferroni’s correction.

We also compared the rates of DNMs for each mutational class. For this analysis, we control for differences in the callable fraction of each trio. We also account for differences in the mutational opportunities in the genome and CGIs by scaling the callable fraction of a given mutational class by the proportion of reference bases, i.e C, T and CpG, in a given genomic context. To test the statistical significance in differences between mutation rates for a specific mutation class in the entire genome and inside CGIs, we used a binomial test from scipy (version 1.7.3) and adjusted the P-values for multiple testing with Bonferroni correction using multipletests from statsmodels (version 0.13.2).

### DNM shared by siblings

In this study, we analyzed 8,565 de novo mutations identified through whole-genome analysis. We catalogued each mutation by its unique chromosomal position and filtered for mutations observed more than once. Subsequently, we examined whether mutations shared between individuals were from siblings or half-siblings, based on parental information. Siblings were defined as individuals with the same parents, while half-siblings were those sharing only one parent All shared mutations in the dataset occurred between either siblings or half-siblings. We identified 79 unique mutations that are shared: 70 of these are common between two individuals, while 9 are shared among three individuals. Notably, only two mutations were found between half-siblings who shared the same father. There are 34 different parental combinations involved in these sibling-shared mutations.

We further compare the mutational spectrums between the 79 shared mutations and the remaining 8,398 unique mutations. This involved calculating the frequency of eight mutation types (C>A, C>G, C>T, T>A, T>C, T>G, C>T in CG context, and mutations in CGIs). For the mutational spectrum, the frequency of all categories, except mutations in CGIs, cumulatively equals 1. We determined the frequency of each mutation type and its 95% confidence interval as follows:

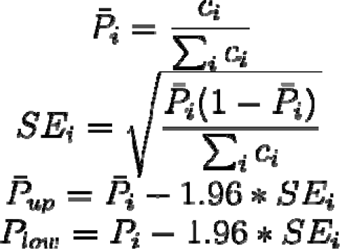

### The impact of mutation and recombination patterns in domestic dogs

We use the branch lengths of the phylogenetic trees to represent the evolutionary rate. We use the whole genome sequences and CGIs region sequences to construct NJ phylogenetic trees and obtain the branch lengths of each branch. We use five species, dog, African wild dog, Arctic fox, polar bear and domestic cat, for the evolutionary rate analysis. The whole genome alignment sequences of the five species come from the HAL alignment of 241 species zoonomia^38^ cactus alignment. The CGIs alignment sequences are extracted from the HAL alignment using the maffilter tool^39^ with the dog as the reference genome. Subsequently, we obtain the species tree with divergence time of the five species from Timetree^40^.

## Data and materials availability

Raw sequence data is available from the GSA (https://ngdc.cncb.ac.cn/gsa/) under accessions CRA004356 CRA002653 CRA002915 and CRA001113; and NCBI (https://www.ncbi.nlm.nih.gov/) under the Bioproject PRJNA1079355.

## Acknowledgments

We thank Sini Karjalainen for technical assistance, Chung-I Wu and Guojie Zhang for helpful comments, and Quankuan Shen for analytical assistance. This project was funded by STI2030-Major Projects 2021ZD0203900 (to G.-D.W.), National Key R&D Program of China 2019YFA0707101 (to G.-D.W.), Jane and Aatos Erkko Foundation (to H.L.), Spring City Plan 2022SCP001 (to G.-D.W.), Yunnan Fundamental Research Projects 202201AV070011, 202201AS070038 and 202305AH340006 (to G.-D.W.), NSFC 32170436 (to G.-D.W.), Novo Nordisk Foundation NNF18OC0031004 and NNF21OC0069105 (to M.H.S.). We downloaded phenotypic information and photos of dog breeds from the website of the American Kennel Club (www.akc.org). This work was supported by the Animal Branch of the Germplasm Bank of Wild Species, Chinese Academy of Sciences (the Large Research Infrastructure Funding). Sequencing data logistics was performed at CSC – IT Center for Science Ltd, Espoo, Finland. (www.csc.fi).

## Author Contributions

G.-D.W., H.L. and M.H.S. contributed to study conceptualization and supervision. G.-D.W. and H.L. design this work. H.L. and J.N. provided samples. S.-J.Z., J.L.M., M.R. analysed data. S.-J.Z., J.L.M., M.R., S.B., T.Z. contributed to visualization. S.B., J.N., N.S., S.H., M.K.H., H.L., G.M.L. and E.A.O. investigate this work. G.-D.W. and S.-J.Z. contributed to project administration. S.-J.Z., J.L.M., M.R., S.B., M.H.S. have completed the original draft. Reviewed and edited by G.-D.W., H.L., M.H.S., E.A.O.

## Competing interests

The authors declare no competing interests.

## Additional information

**Supplementary Information** is available for this paper.

**Correspondence and requests for materials** should be addressed to Guo-Dong Wang.

**Extended Data Fig. 1.**
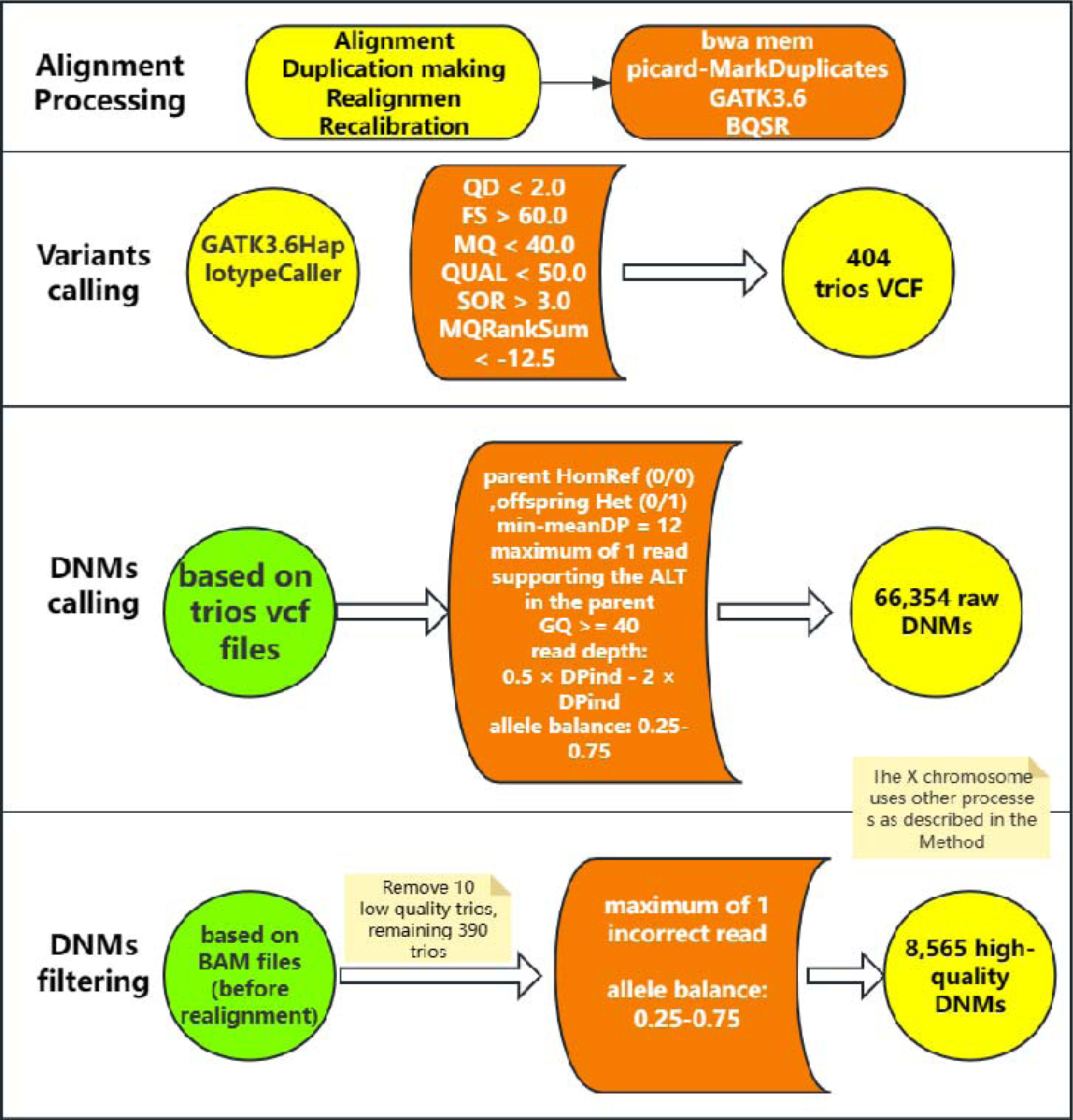
Flowchart for the identification of de novo mutations.

**Extended Data Fig. 2.**
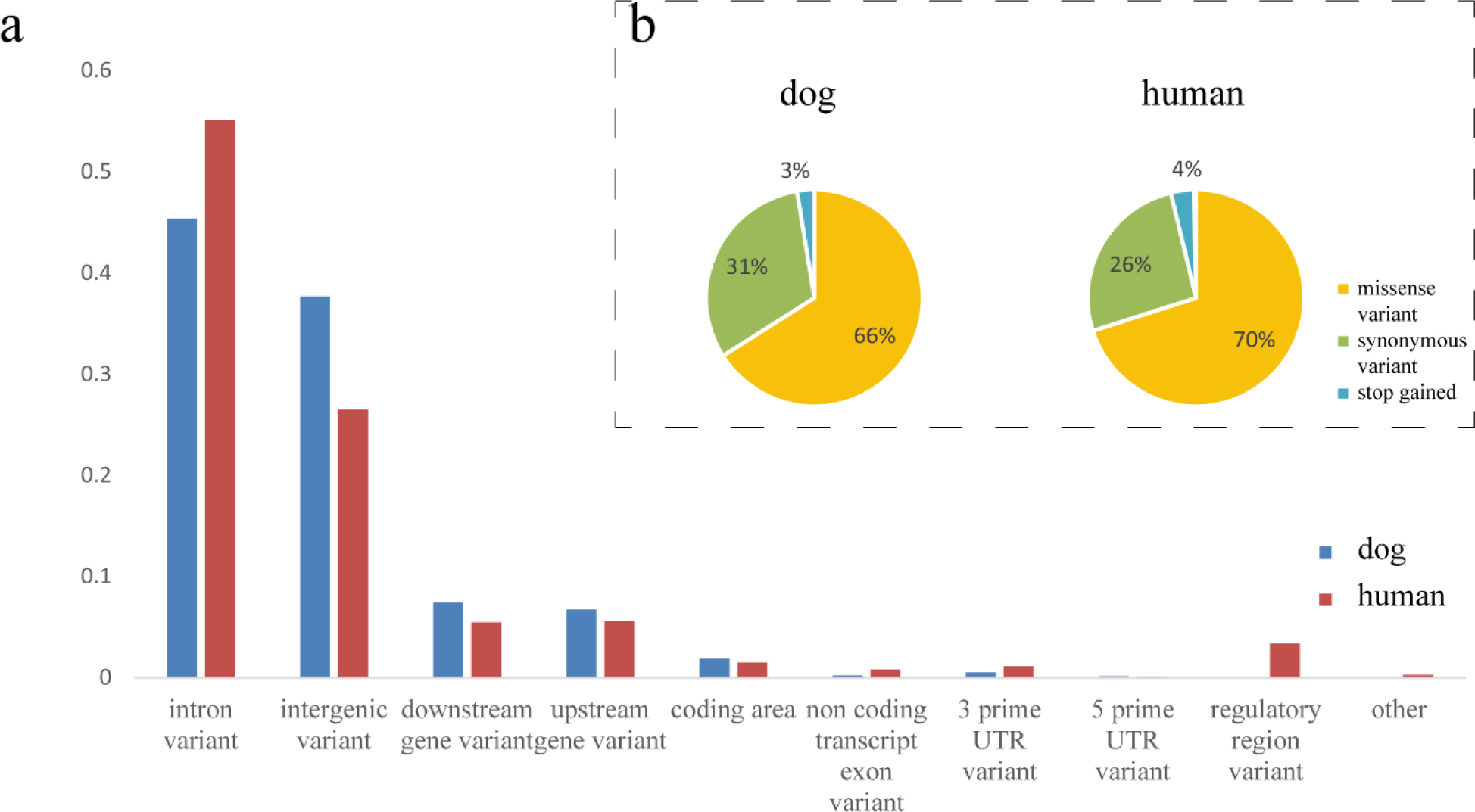
Variant type difference of de novo mutations between dog and human. (a) Comparison in all de novo mutations. (b) Comparison in coding region mutations.

**Extended Data Fig. 3.**
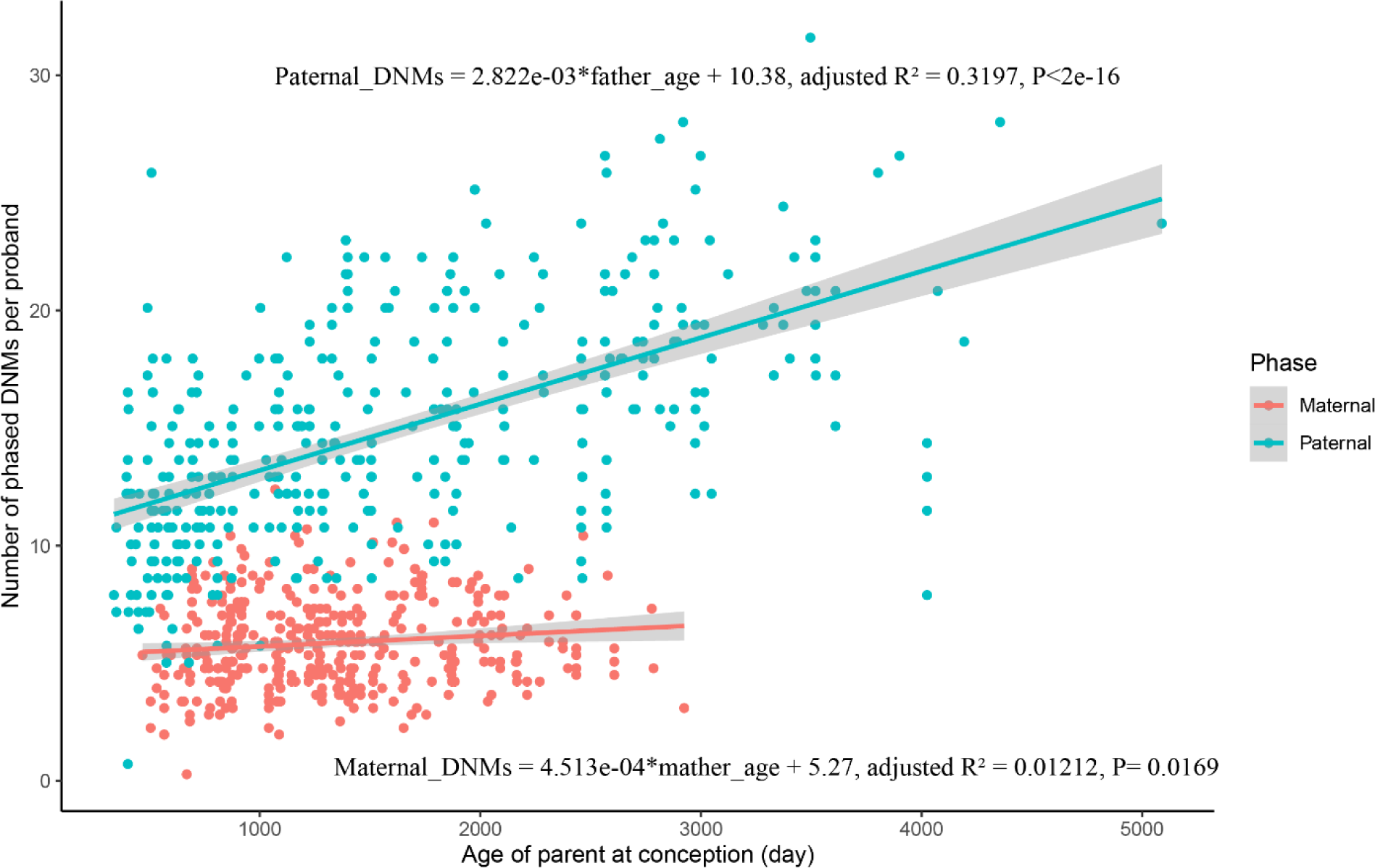
Phased DNMs as a function of the parent’s age at conception.

**Extended Data Fig. 4.**
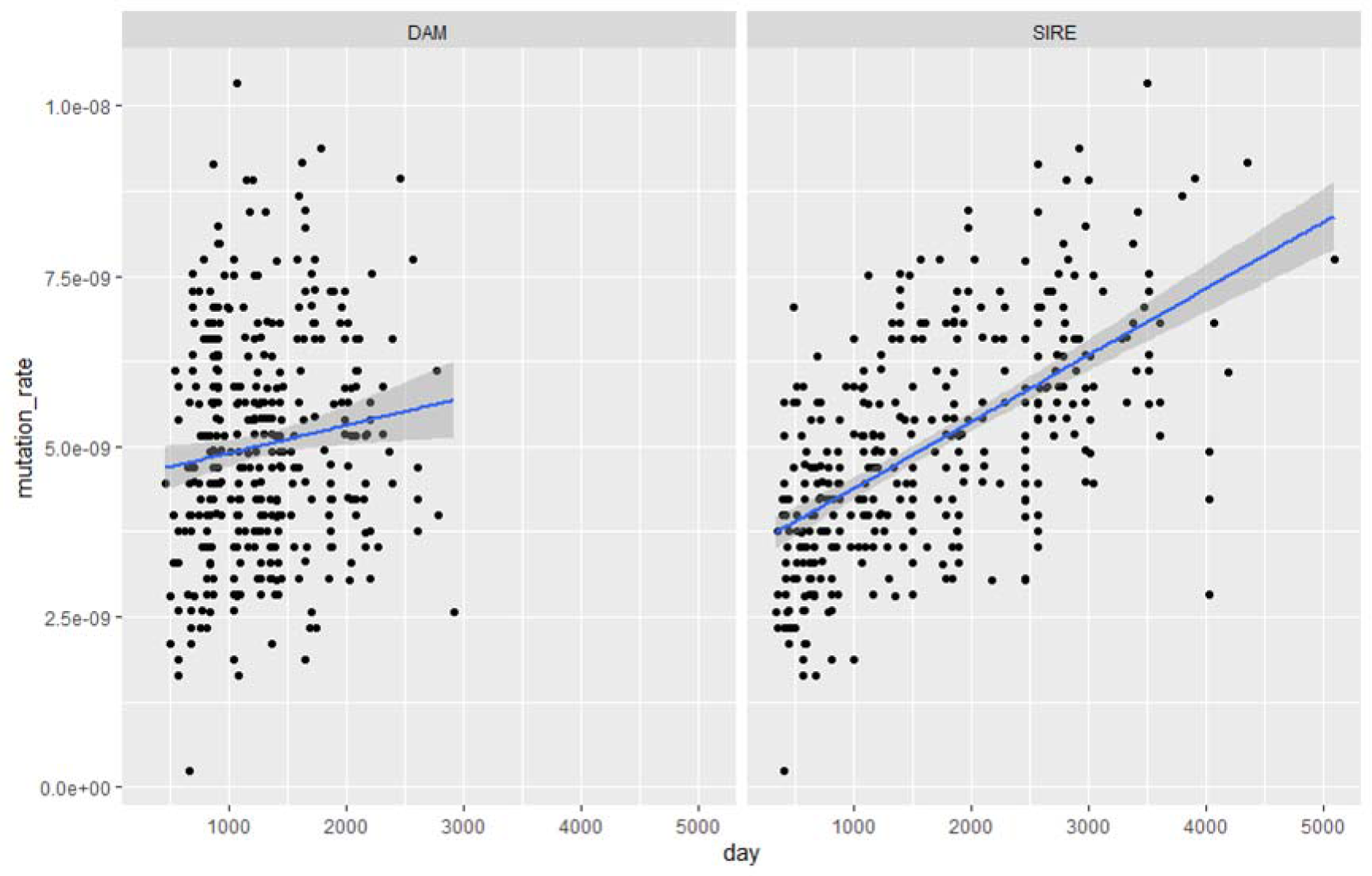
Parent’s age as a function of the mutation rate (per generation).

**Extended Data Fig. 5.**
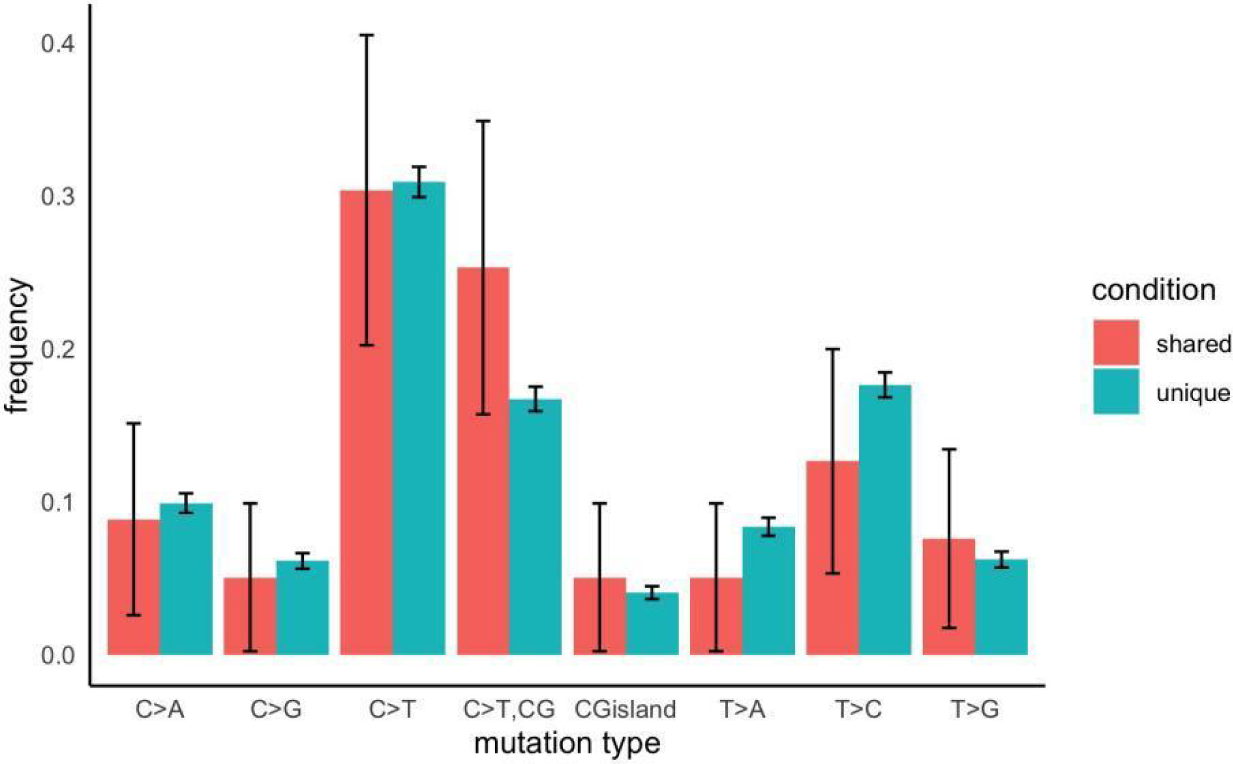
Mutational spectrum in shared mutations and non-shared mutations.

**Extended Data Table 1.**
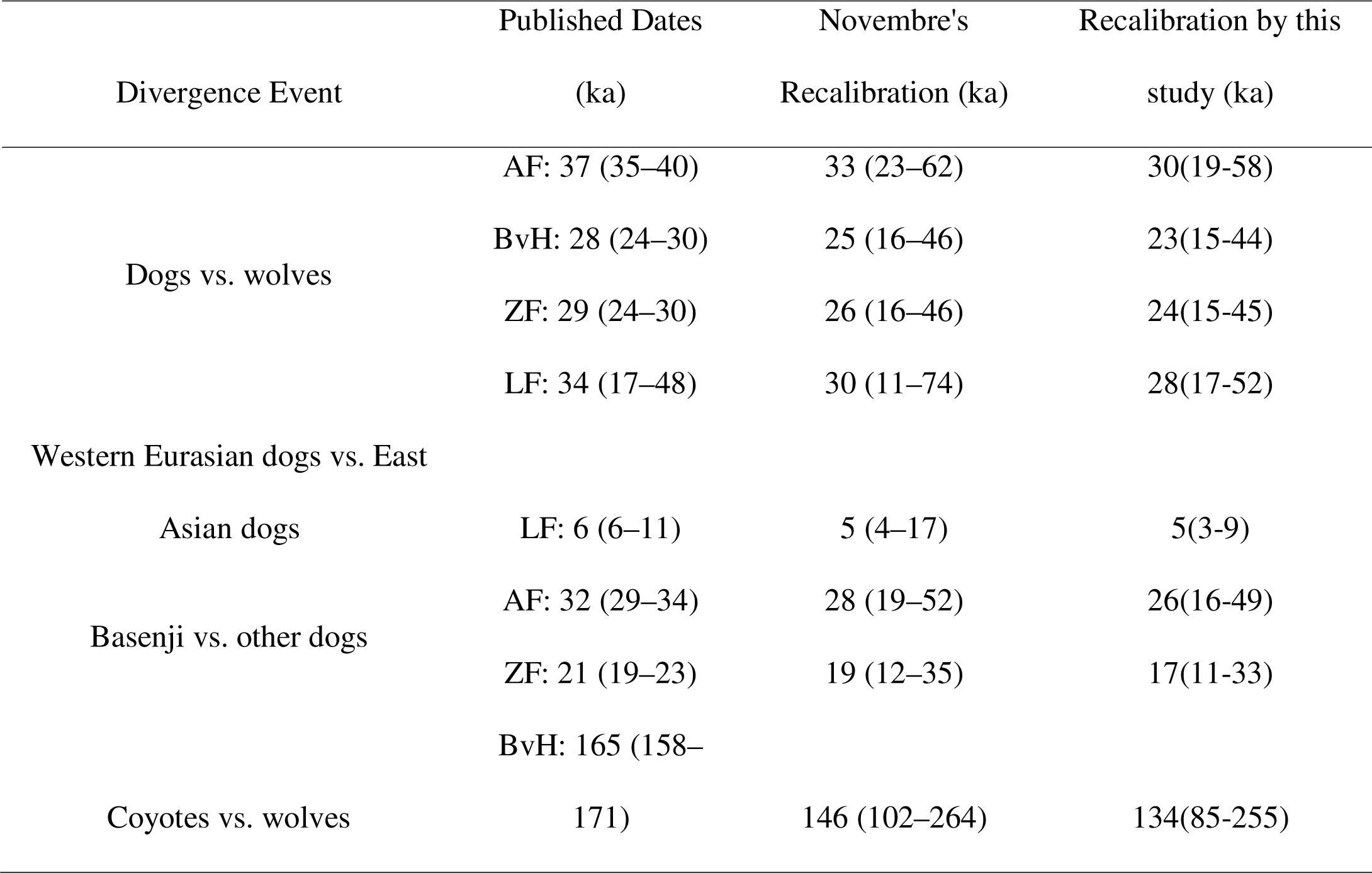
Recalibration of estimated divergence times in canid.

## Notes

### Competing Interest Statement

The authors have declared no competing interest.

