## Supplemental Information for "Determinants of de novo mutations in extended pedigrees of 43 dog breeds"

**Supplementary information 1: Sample collection**

The dogs were selected from the Finnish dog biobank, containing nearly 80 000 canine DNA samples donated by private dog owners. All dogs in the study originate from Finland. We assembled trios and additional siblings for sequencing from two different types of pedigree cohorts: a pedigree cohort with 3-5 generations, including both parents and littermates, and a sire cohort, in which we had sires that had up to 5 litters with different bitches over ~10 years period (Figure 1, Fig. S1, Data S1). We maximized the number of different breeds with different characteristics, such as size and appearance, for the study from the samples available in the biobank. EDTA blood samples were stored at -20°C until genomic DNA was extracted using a semi-automated Chemagen extraction robot (PerkinElmer Chemagen Technologie GmbH). DNA concentration was determined either with NanoDrop ND-1000 UV/Vis Spectrophotometer, Qubit 3.0 Fluorometer (Thermo Fisher Scientific Inc.) or DeNovix DS-11 Spectrophotometer (DeNovix Inc., Wilmington, Delaware, USA).

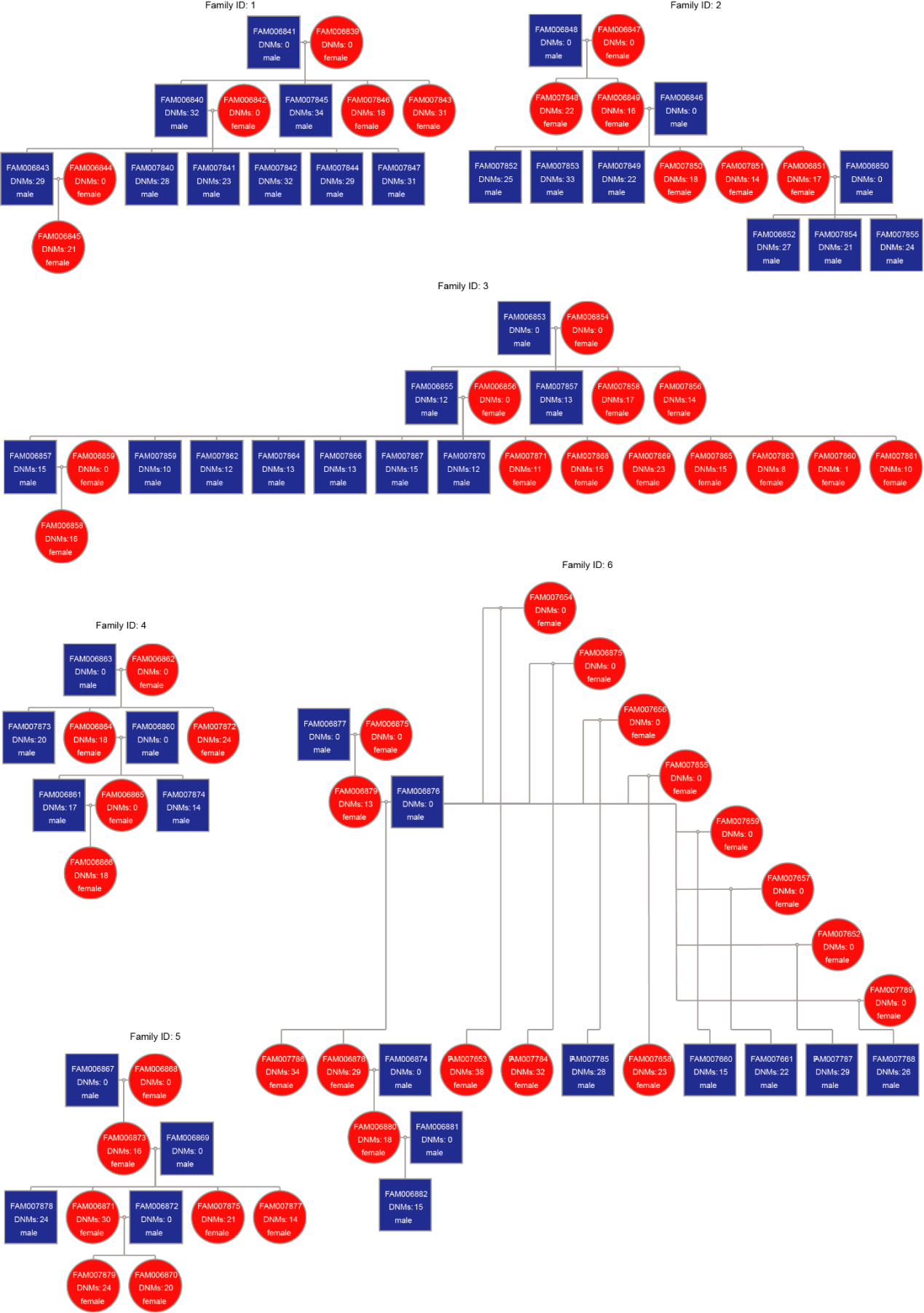

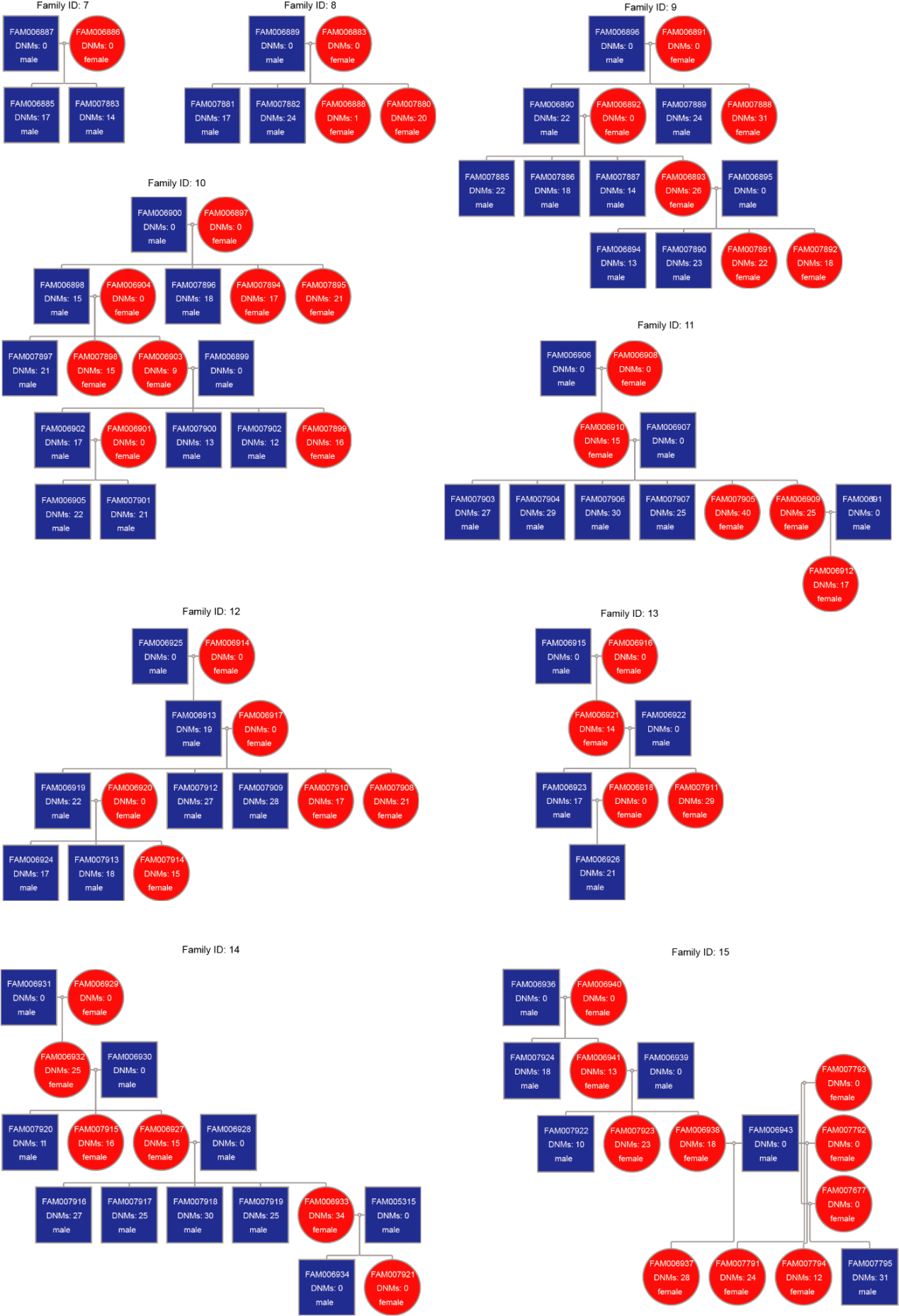

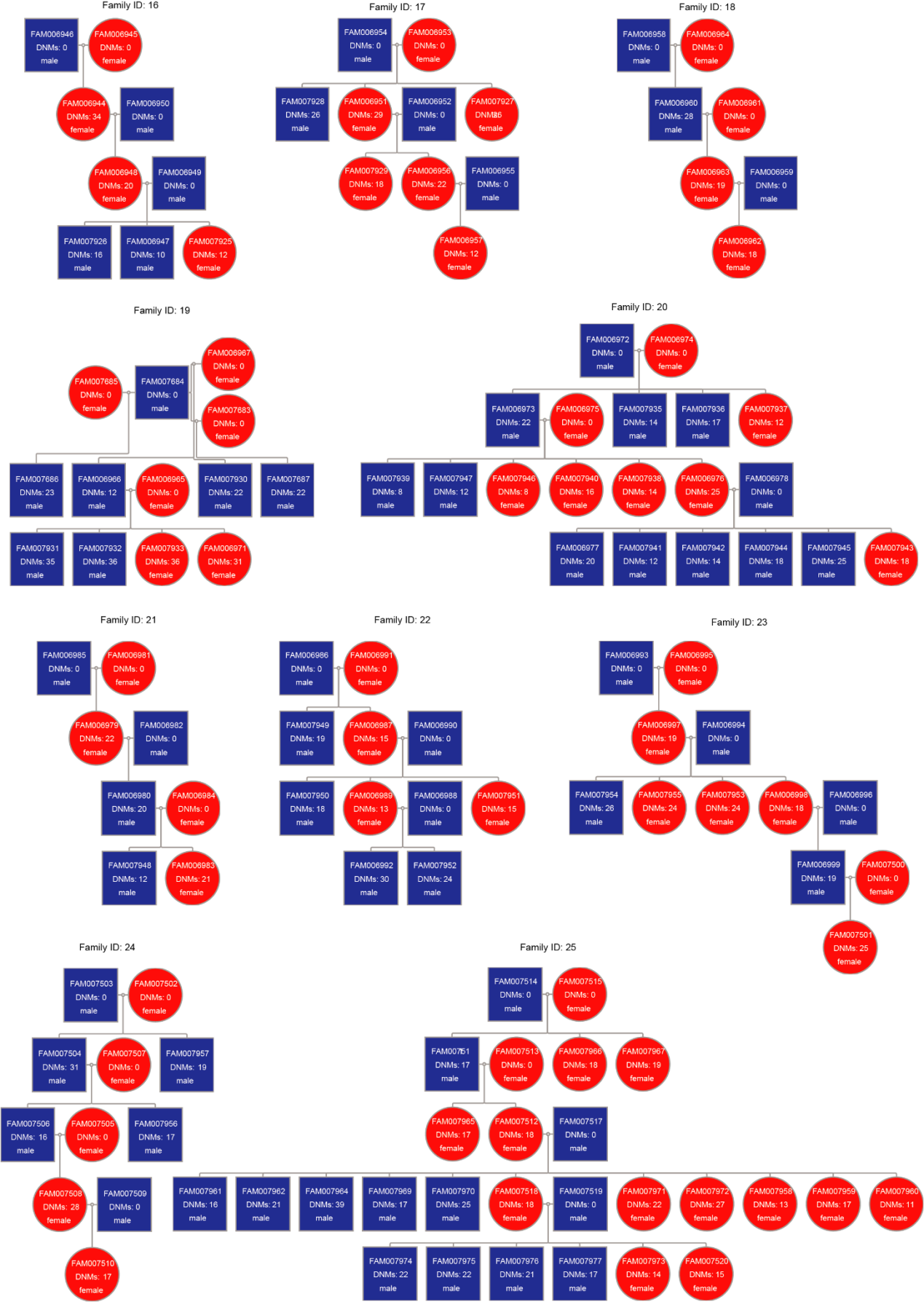

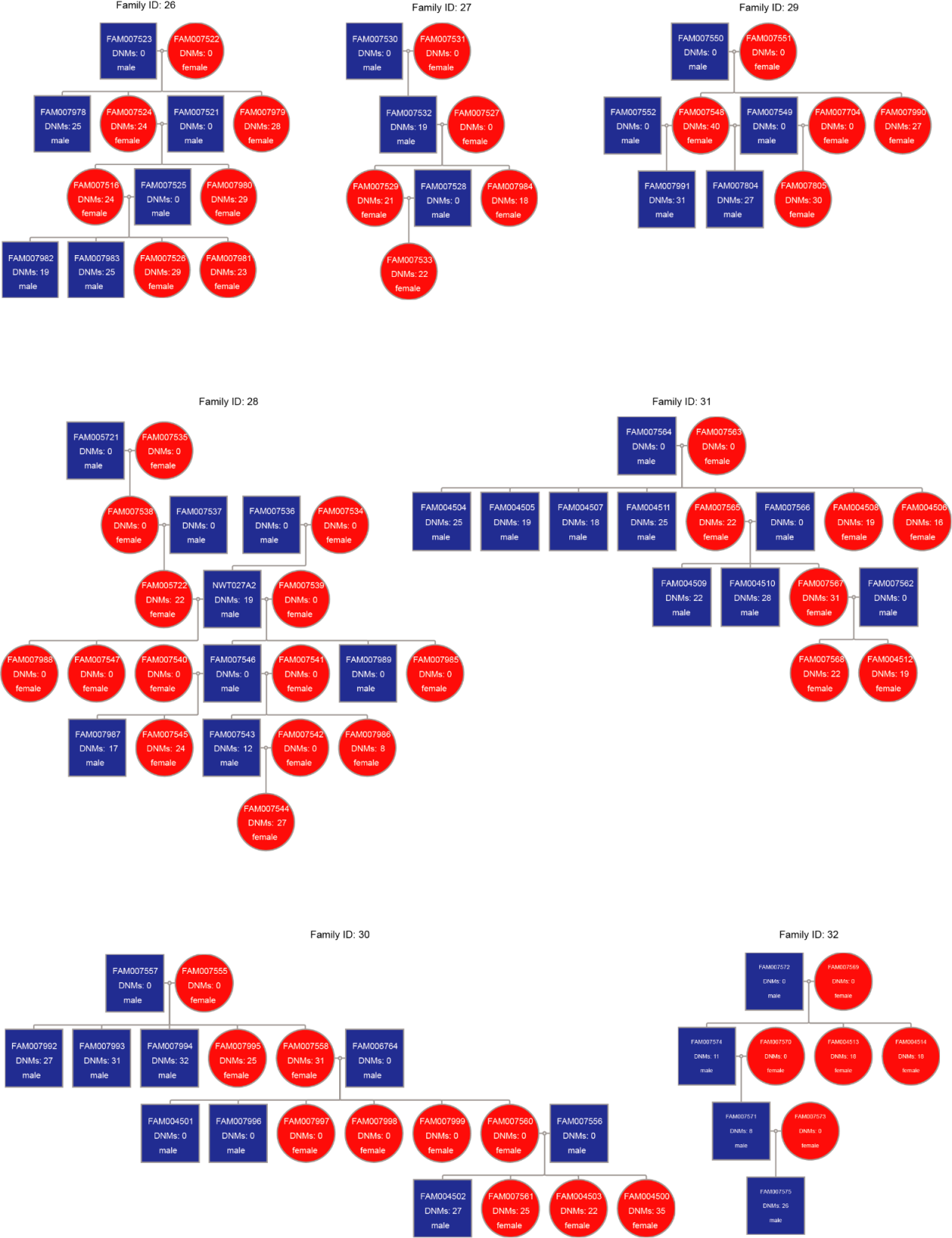

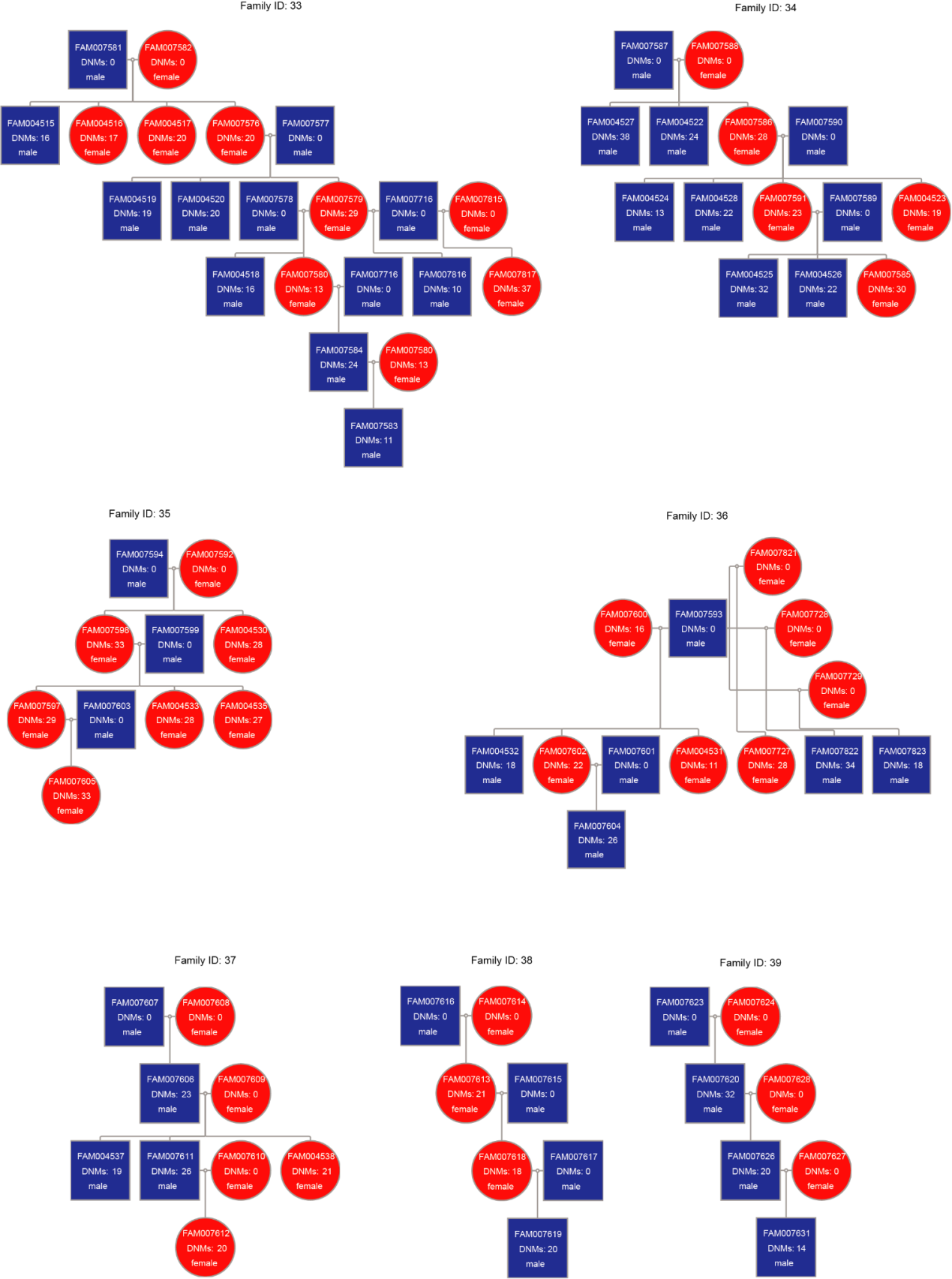

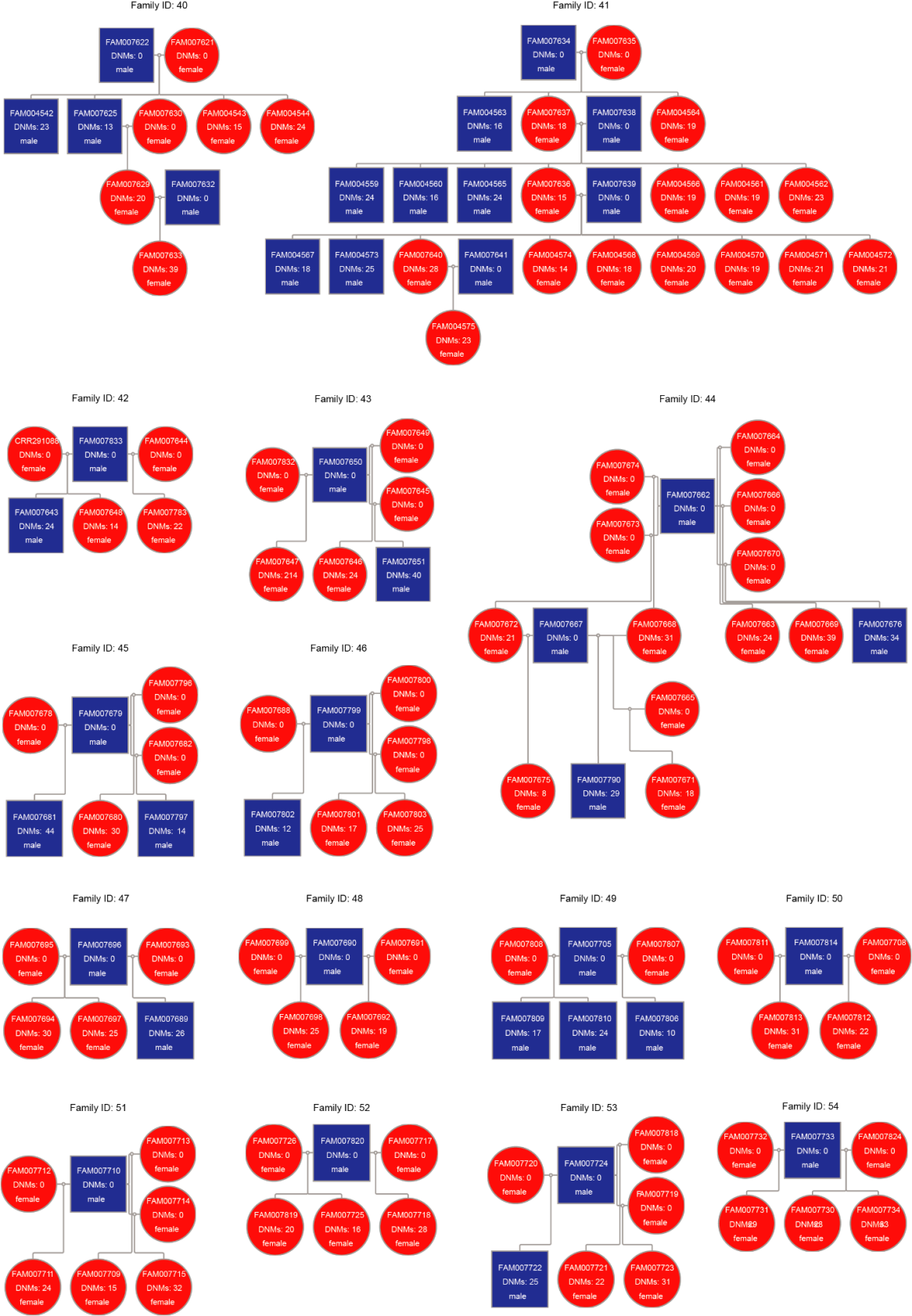

**Fig. S1.**

Pedigree chart of 54 domestic dog families.

**Supplementary information 2: Mutational landscape and enrichment of hypermutation genes**

We performed the gene annotation using the Variant Effect Predictor (VEP)^1^ in the Ensembl. We use the "maftools" package^2^ in R to visualize the results of DNMs annotation, and then get mutational landscape. 8,312 dog autosomal DNMs have been found in 3,126 genes, but just 54 genes (1.71%) had more than 3 DNMs in their gene region (Fig. S2, Data S2). We called the gene with unusually large numbers of DNMs as Hypermutation gene. We found 106 hypermutation genes in human DNMs dataset using the same method, which have over 60 DNMs (Fig. S3, Data S3). Enrichment analysis is performed using “g:GOSt” module in g:Profiler^3^. The hypermutated genes of human and dog were both enriched in terms related to synapse and nervous system development(Data S4 and Data S5). Many of these genes play very important roles in neurons and synapses.

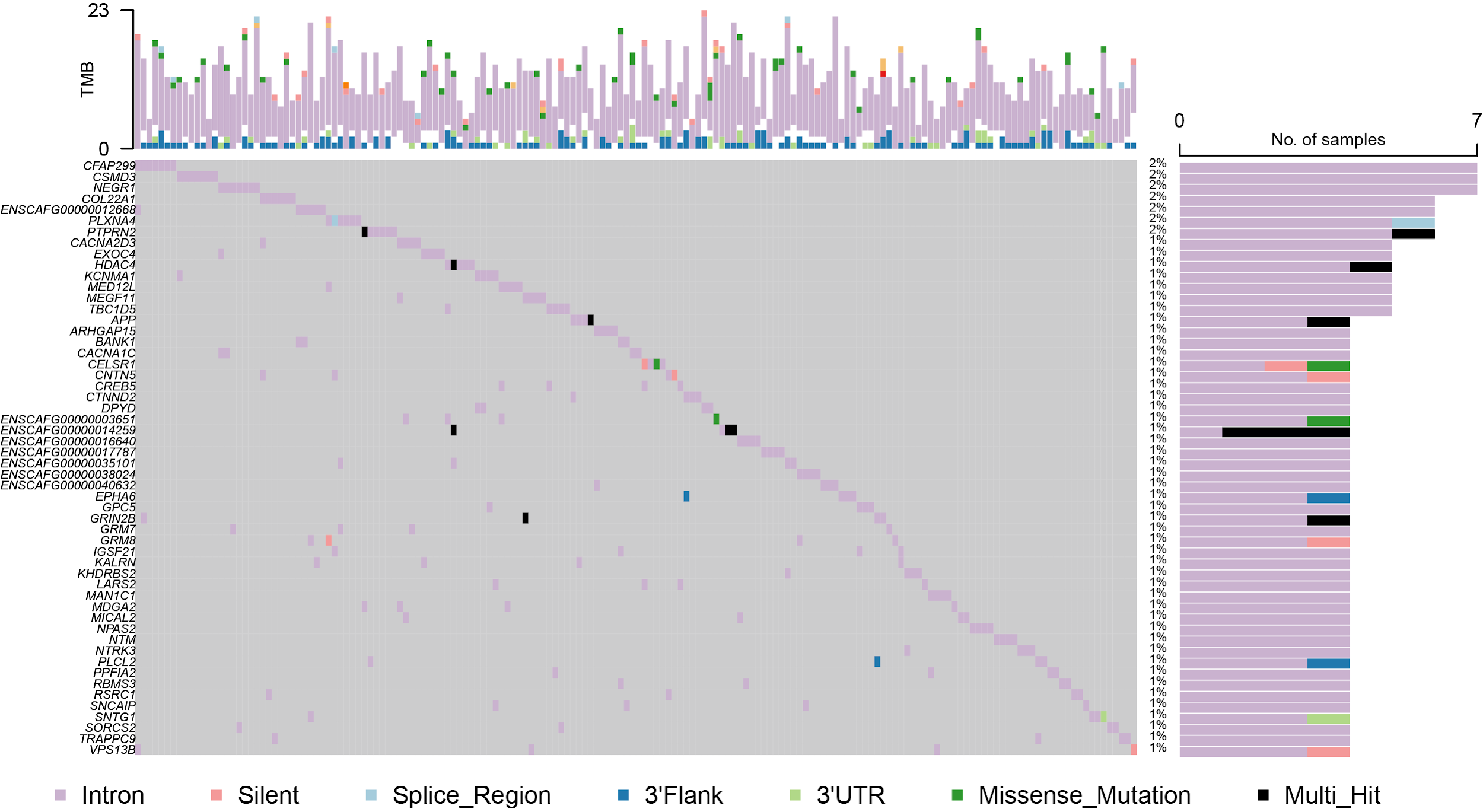

**Fig. S2.**

Mutational landscape of DNMs (except for the DNMs in intergenic regions). Rows are genes and columns are samples, the right panel shows number of samples, the upper panel shows the number of DNMs in each sample, and different mutation types are represented by different colors.

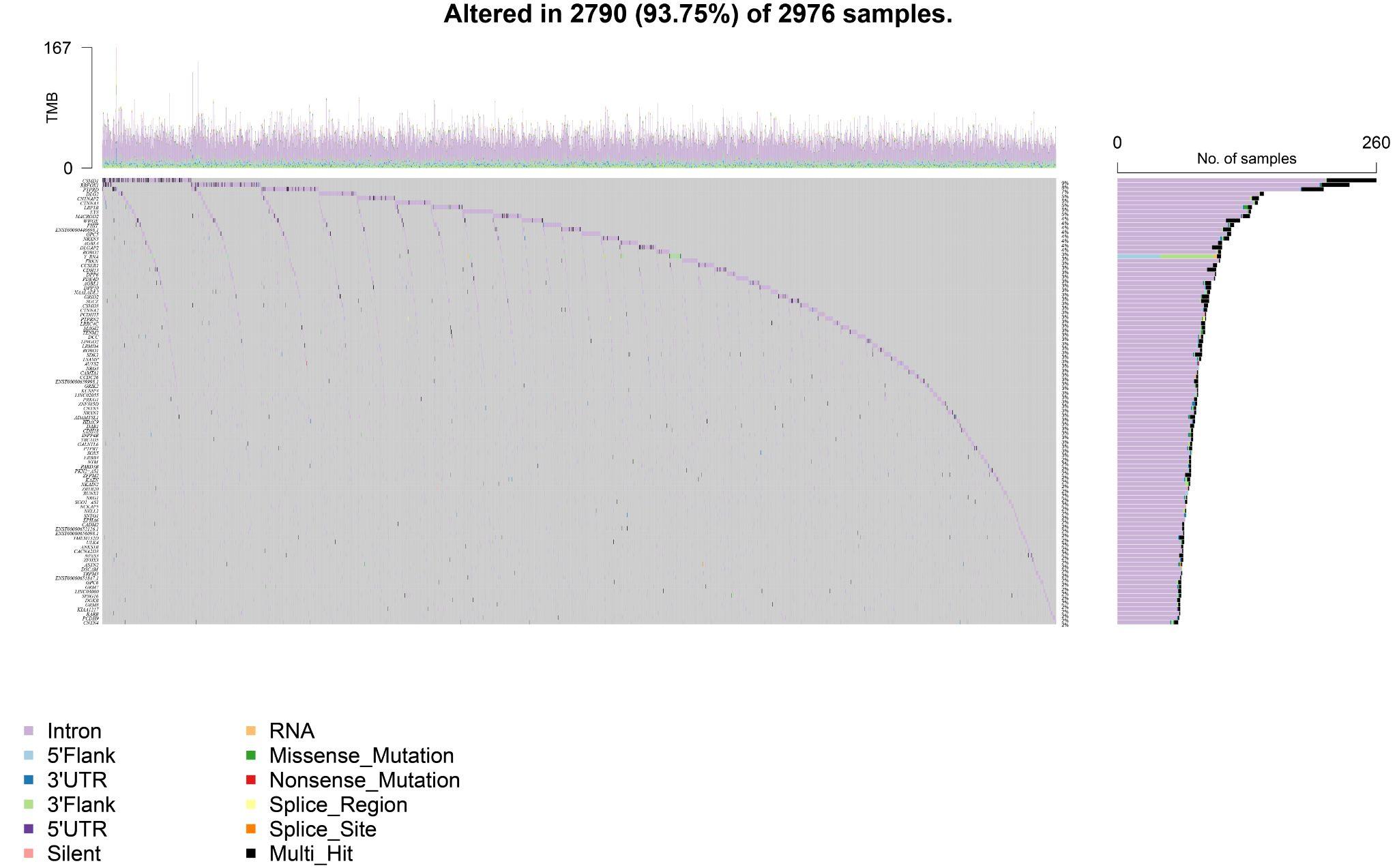

**Fig. S3.**

Mutational landscape of human DNMs.

**Supplementary information 3: Correlation analysis of mutation rate**

3.1 Breed effect on mutation rate

We modelled mutation rates for all trios as a function of breed using linear regression, accounting for paternal and maternal ages, with the lm function in R (Fig. S4, S5). We note that the structure of our dataset can bias the results on the breed effect on mutation rates, since trios are organized in litters and the number of trios per litter is different for each breed. Since multiple trios from the same breed are expected to have the same mutation rate, we accounted for this possible bias by using the average mutation rates across litters instead of trios. We also ran an ANOVA analysis on per trio and per litter mutation rates as a function of parental ages and breed with the aov function in R (Fig. 2a).

3.2 Phenotype effect on mutation rates

We also collected 21 breeds’ characteristics from American Kennel Club (AKC) and quantified them as phenotypic values of breeds (Table S1). We then modelled per litter mutation rates as a function of these phenotypic traits, accounting for paternal and maternal ages, with the lm function in R. (Table S7).

Randomly select one individual for each breed as the sample set for phylogenetic analysis. And then, The NJ phylogenetic tree was built by SNPhylo^4^. The pictures of each breed dog are from AKC (www.akc.org).

3.3 Germline mutation rate by genome regions

We assign the whole genome into 4 different regions including autosomal regions, CGIs, PAR, and X chromosome unique regions. The autosomal regions contain all autosomes. The CGIs correspond to CGIs on autosomes based on CanFam3.1 annotation. Position 1:680000 from X chromosome is retrieved as the PAR in our analysis and the rest of X chromosome are noted as the unique part of X chromosome.

Table S1. The result of 21 phenotype effect on per litter mutation rates

| variable | pvalue |
| --- | --- |
| DROOLING_LEVEL | 0.1323 |
| COAT_GROOMING_FREQUENCY | 0.3892 |
| OPENNESS_TO_STRANGERS | 0.4183 |
| ADAPTABILITY_LEVEL | 0.4622 |
| AFFECTIONATE_WITH_FAMILY | 0.4993 |
| SHEDDING_LEVEL | 0.5159 |
| Litter_size | 0.5617 |
| ENERGY_LEVEL | 0.6077 |
| GOOD_WITH_YOUNG_CHILDREN | 0.6213 |
| TRAINABILITY_LEVEL | 0.6764 |
| PLAYFULNESS_LEVEL | 0.7099 |
| Height_male | 0.7198 |
| WATCHDOG | 0.7282 |
| GOOD_WITH_OTHER_DOGS | 0.7286 |
| Lifespan | 0.7646 |
| MENTAL_STIMULATION_NEEDS | 0.7723 |
| Height_female | 0.7784 |
| Size | 0.7956 |
| BARKING_LEVEL | 0.8351 |
| Weight_male | 0.8445 |
| Weight_female | 0.8682 |

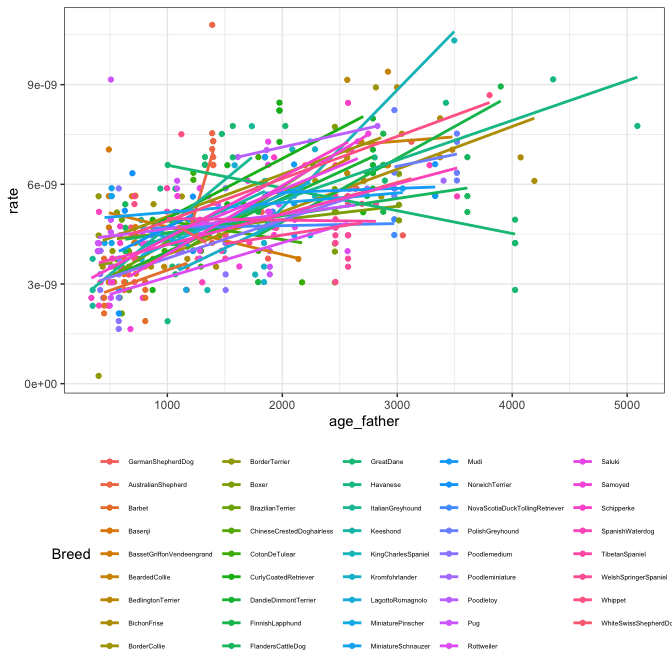

**Fig. S4.**

Correlation analysis between mutation rate (per trio) and breed.

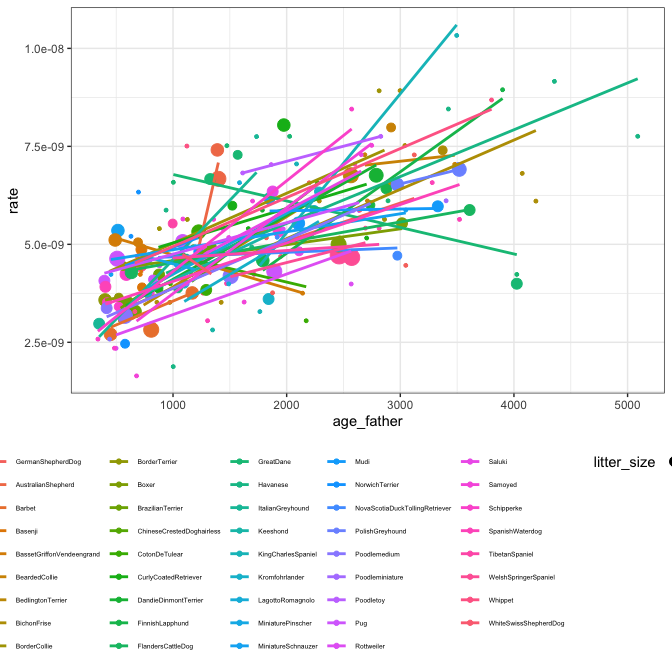

**Fig. S5.**

Correlation analysis between mutation rate (per litter) and breed.

**Supplementary information 4:** **Parental age effect on germline mutation rates**

4.1. Linear regression on germline mutation rates

We modelled mutation rates for each region as a function of paternal and maternal ages using linear regression with the lm function in R.

We also modelled mutation rates for each breed as a function of paternal and maternal ages with the lm function in R.

4.2. Poisson regression on upscaled DNMs

For this analysis we only considered trios with phased DNMs, thus retaining 347 dog trios. Similar to previous work (Riera et al. X), we modeled the phase-specific accumulation DNMs (
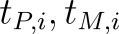
) with Poisson regression. We define lambda (
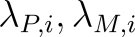
) as a function of paternal and maternal age at conception (
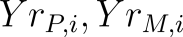
) and account for differences in the callable fraction for each trio.

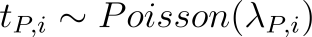

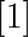

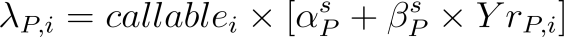

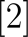

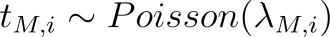

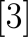

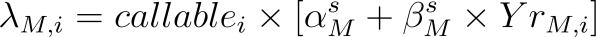

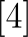

With s being the species in model X, i.e dog and human, and s being the size group in dogs and species in humans in model X.

For simplification purposes, we scaled
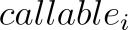
 by 10^-10^, thus moving the range of possible values for
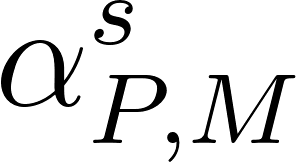
 and
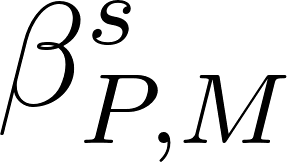
 away from zero. After fitting the model, we scaled the obtained estimates back to the original range.

We define
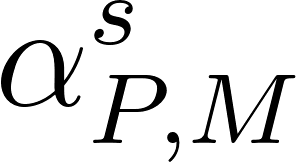
 with a HalfNormal distribution, using a shared hyperprior across species, and
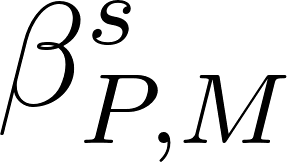
 with an Exponential distribution, parametrized as following:

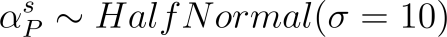

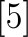

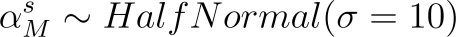

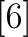

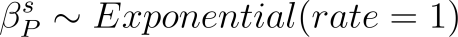

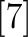

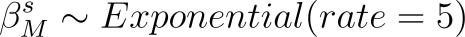

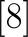

4.3. Germline yearly mutation rates.

We calculated germline yearly mutation rates using the predicted mutation rates from our Poisson regression fit model at the average generation time for each sex and species. Following a previous model^5^, we accounted for differences in the generation time of males and females (
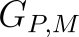
<https://www.codecogs.com/eqnedit.php?latex=G_%7BP%2CM%7D#0>):

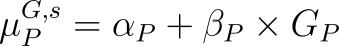
<https://www.codecogs.com/eqnedit.php?latex=%5Cmu_%7BP%7D%5E%7BG%2Cs%7D%20%20%3D%20%5Calpha_%7BP%7D%20%2B%20%5Cbeta_%7BP%7D%20%5Ctimes%20G_%7BP%7D%20#0>
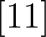
<https://www.codecogs.com/eqnedit.php?latex=%5B11%5D#0>

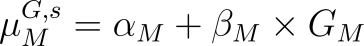
<https://www.codecogs.com/eqnedit.php?latex=%5Cmu_%7BM%7D%5E%7BG%2Cs%7D%20%20%3D%20%5Calpha_%7BM%7D%20%2B%20%5Cbeta_%7BM%7D%20%5Ctimes%20G_%7BM%7D%20#0>
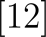
<https://www.codecogs.com/eqnedit.php?latex=%5B12%5D#0>

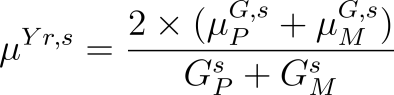
<https://www.codecogs.com/eqnedit.php?latex=%20%5Cmu%5E%7BYr%2Cs%7D%20%3D%20%5Cfrac%7B2%20%5Ctimes%20(%5Cmu%5E%7BG%2Cs%7D_%7BP%7D%20%2B%20%5Cmu%5E%7BG%2Cs%7D_%7BM%7D)%7D%7BG_%7BP%7D%5E%7Bs%7D%20%2B%20G_%7BM%7D%5E%7Bs%7D%20%7D%20#0>
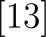
<https://www.codecogs.com/eqnedit.php?latex=%5B13%5D#0>

4.4. Downsampling

We did 100 random down-samples of the human DNM dataset to match the number of dog trios with phased DNMs (N = 347) and the average number of DNMs per trio in dogs by dividing the number of DNMs in a given human trio by a factor of 3. For each human down-sample and the dog dataset we ran a Poisson regression on mutation counts as a function of paternal age using the glm function on R (version 4.2.2). We calculated McFadden’s R^2^ as 1 minus the ratio of the deviance to the null deviance of the model fit (Fig. S6, S7).

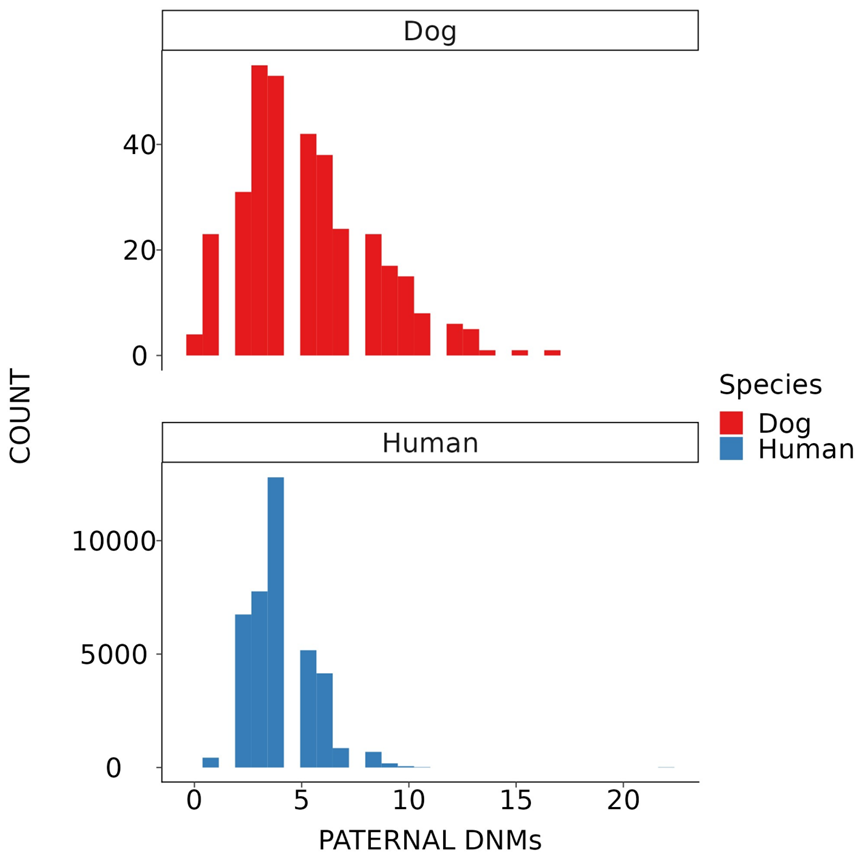

**Fig. S6.**

Distribution of paternal DNM counts in dogs and humans, after down-sampling the human dataset to match the expected number of DNMs found in dogs.

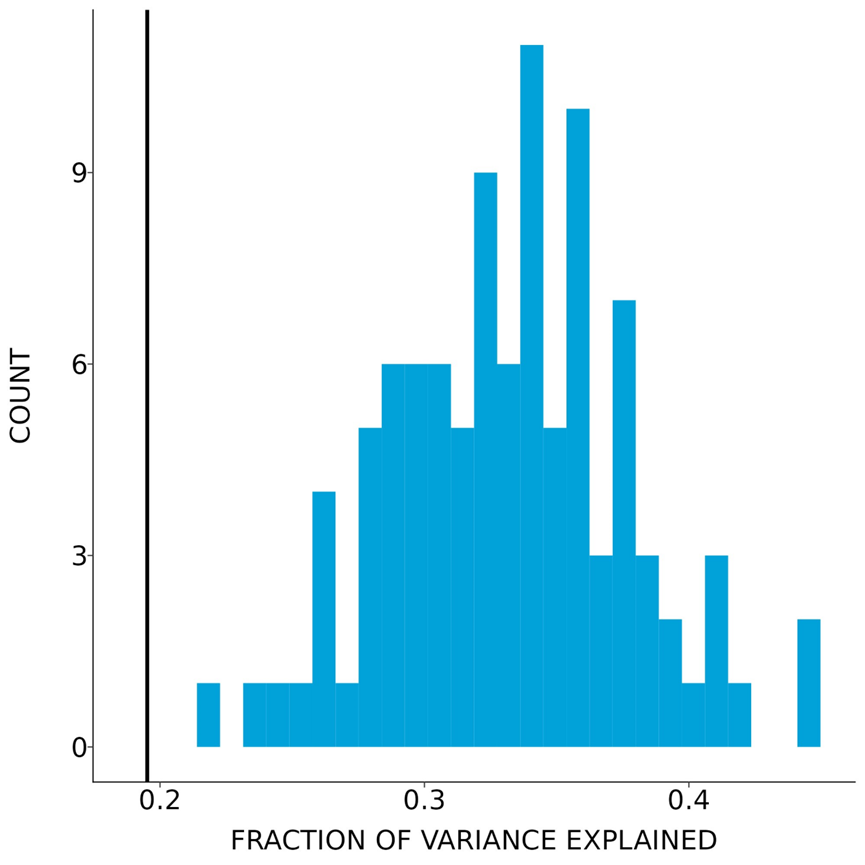

**Fig. S7.**

Distribution of McFadden’s R2 values in blue obtained from Poisson regression on paternal germline DNMs as a function of the age of the father in 100 random down samples of human DNMs to the average number of DNMs in dogs. The value for McFadden’s R2 obtained in dogs is shown in black.
